# A novel *Phytophthora sojae* effector PsFYVE1 modulates transcription and alternative splicing of immunity related genes by targeting host RZ-1A protein

**DOI:** 10.1101/2021.12.02.470886

**Authors:** Xinyu Lu, Zitong Yang, Wen Song, Jierui Si, Zhiyuan Yin, Maofeng Jing, Danyu Shen, Daolong Dou

**Affiliations:** Key Laboratory of Plant Immunity, Academy for Advanced Interdisciplinary Studies, College of Plant Protection, Nanjing Agricultural University, Nanjing 210095, China

## Abstract

Oomycete pathogens secrete many effectors to manipulate plant immunity and promote infection. However, relatively few effector types have been well characterized. In this study, members of a FYVE domain-containing protein family that is highly expanded in oomycetes were systematically identified, and one secreted protein, PsFYVE1, was selected for further study. PsFYVE1 enhanced *Phytophthora* infection in *Nicotiana benthamiana* and was necessary for *P. sojae* virulence. The FYVE domain of PsFYVE1 had PI3P-binding activity that depended on four conservative amino acid residues. Furthermore, PsFYVE1 targeted RNA-binding proteins RZ-1A/1B/1C in *N. benthamiana* and soybean, and silencing of *NbRZ-1A*/*1B*/*1C* genes attenuates plant immunity. NbRZ-1A was associated with spliceosome that included three important components, NbGRP7, NbGRP8, and NbU1-70K. Notably, PsFYVE1 could disrupt NbRZ-1A–NbGRP7 interaction. RNA-seq and subsequent experimental analysis demonstrated that PsFYVE1 and NbRZ-1A not only co-regulated transcription of *NbHCT*, *NbEIN2*, and *NbSUS4* genes but also modulated pre-mRNA alternative splicing (AS) of the *NbNSL1* gene, which participated in plant immunity. Collectively, these findings indicate that the FYVE domain-containing protein family includes potential new effector types and also highlight that plant pathogen effectors can regulate plant immunity related genes at both transcription and AS levels to promote disease.

**Author summary:** Many plant pathogenic oomycetes secrete effector proteins into plants to facilitate infection. Discovering potential repertoire of novel effectors and corresponding molecular mechanisms are major themes in the study of oomycete–plant interactions. Here, we characterized a FYVE domain-containing protein (PsFYVE1) in *P. sojae*. PsFYVE1 carries a functional secretory signal peptide and is a virulence-essential effector for *P. sojae* infection. We demonstrated that PsFYVE1 interacted with a class of plant RNA-binding proteins, including soybean GmRZ-1A/1B/1C and *N. benthamiana* NbRZ-1A/1B/1C. Silencing of *NbRZ-1A*/*1B*/*1C* proteins increased *Phytophthora* infection and suppressed plant defense. Furthermore, NbRZ-1A interacted with the spliceosome components, and PsFYVE1 disrupted association between NbRZ-1A and spliceosome component NbGRP7. We examined the global transcription and alternative splicing (AS) changes regulated by PsFYVE1 and NbRZ-1A, which indicated that PsFYVE1 and NbRZ-1A co-regulated transcription and pre-mRNA AS of immunity-related genes. Thus, this study identifies a novel virulence-related effector from *P. sojae* and a class of positive regulators of plant immunity, and reveals a detailed mechanism of effector-medicated transcription and AS regulation during pathogen–plant interactions.

## Introduction

Oomycetes such as *Phytophthora* spp. include a wide variety of plant pathogens that cause devastating diseases on a wide range of crops [1]. During infection, *Phytophthora* pathogens secrete several types of effector proteins into hosts to subvert plant defense responses [2]. Apoplastic effectors are one type and include elicitins, necrosis- and ethylene-inducing peptide 1 (NEP1)-like proteins (NLPs), glycoside hydrolase 12 proteins (GH12), and cellulose-binding elicitor lectin (CBEL), which contain the signal peptides and are secreted into the host extracellular space during interaction [3, 4]. Other type is cytoplastic effectors, such as RXLR and Crinkler (CRN) that contain an N-terminal signal peptide and are delivered into plant cells [5]. Recently, some specific effectors are also reported in oomycetes; for example, the unconventionally secreted effectors [6], the *Albugo* CHXC effectors [7], and the small open reading frame-encoded polypeptides [8]. However, investigations on virulence strategies of oomycete pathogens tend to be restricted to conventional effector types. Furthermore, prediction of oomycete secreted effectors by common sequence features will miss bona fide effectors that do not carry such signatures [9]. RXLR effectors have been studied extensively, because they can target plant proteins to interfere with host immunity through sophisticated pathogenicity mechanisms, in turn, some of them may be recognized by plant NLR-mediated immune responses [10–12]. However, how RXLR effectors enters host cells is subject to debate. Kale *et al*. propose that the RXLR domain binds to phosphatidylinositol 3-phosphate (PI3P) to mediate translocation of effector proteins into plant cells and to increase their stability *in planta* [13]. Nevertheless, Gan *et al*. does not demonstrate that flax rust RXLR effectors AvrM and AvrL567 require PI3P binding to internalize into plant cells [14]. According to Yaeno *et al*., AVR3a binds to PI3P via the effector domain, not the RXLR domain, although PI3P-binding ability is essential for its accumulation inside host cells [15].

The phosphoinositide PI3P associates with endosomal membranes to regulate protein trafficking and endocytic pathways, autophagy, and cytokinesis in various eukaryotes [16]. During interactions between plants and pathogens, phosphoinositides are crucial for recognition and membrane trafficking to hijack host cellular processes. For example, during *Colletotrichum higginsianum* infection, both PI(4,5)P_2_ and PI4P are at the extra-invasive hyphal membrane (EIHM), a plant cell-derived membrane that surrounds the hemibiotrophic hyphae [17]. In the Arabidopsis-powdery mildew interaction, only PI(4,5)P_2_ is recruited to the extrahaustorial membrane (EHM) derived from the host plasma membrane (PM). Deletion of *PIP5K* genes responsible for PI(4,5)P_2_ biosynthesis prevents susceptible responses and colonization of powdery mildew in leaf tissues [18, 19]. In our previous study, *Phytophthora*-derived PI3P is exposed at the surface of infection hyphae and plant cell plasma membranes to aid the actions of RXLR effectors [20]. Thus, the compound can be used to guide anti-microbial peptides produced by plant cells to accumulate specially at pathogen tissues to achieve high disease resistance [21], further suggesting important roles of PI3P in *Phytophthora* infection.

The PI3P-binding FYVE domain, is named after the first four FYVE proteins, Fab1p, YOTB, Vac1p, and EEA1. The FYVE domain contains conserved coordinating residues: the N-terminal WxxD, the central R(R/K)HHCR, and C-terminal R(V/I)C residues, which are the principal sites of compact binding with PI3P [16, 22]. The FYVE domain is widely distributed in eukaryotes. According to previous studies, 27 proteins contain the FYVE domain in humans, 5 in yeast, and 15 in *Caenorhabditis elegans* [16]. The FYVE proteins regulate membrane trafficking, receptor signaling, and cytoskeletal dynamics by specific binding to PI3P [23]. The human FYVE protein SARA (Smad anchor for receptor activation) initiates a signaling cascade by recruiting intracellular signaling mediators to receptors [24]. Recently, one *P. sojae* FYVE protein PsFP1 is found to play an important role in mycelial growth, pathogenicity and oxidative stress response [25]. However, the distribution and biological functions of other FYVE proteins in oomycete pathogens are poorly known.

Transcriptional regulation and alternative pre-mRNA splicing are important in plant adaptations to the environment [26]. The processes provide an important RNA-based layer of protein diversity regulation in development and stress response [27]. For example, a plant peptide hormone and its receptor complex RALF1-FER modulate dynamic alternative splicing (AS) in cold response and ABA response by interacting with glycine-rich RNA-binding protein 7 (GRP7) [28]. Moreover, evidence is emerging that various biotic (viral and oomycete pathogens) and abiotic stresses (heat and cold) can trigger feedback on transcriptions and AS. For example, *P. sojae* effector PsCRN108 inhibits expression of plant heat shock protein (Hsp)-encoding genes by directly targeting their promoters to suppress plant *HSP* gene transcription [29]. In addition, an avirulent effector PsAvr3c derived from *P. sojae* reprograms host pre-mRNA splicing by binding to serine/lysine/arginine-rich proteins SKRPs associated with spliceosome complex [30]. These observations indicate that gene transcription and AS are important components of host transcriptome reprogramming in response to pathogens infection.

Sequence-specific glycine-rich RNA-binding proteins (GRPs) are critical for RNA processing in various organisms and have been intensively investigated in the plant kingdoms. Plant GRPs usually contain a canonical RNA recognition motif (RRM) or a cold-shock domain (CSD) at the N-terminus and a glycine-rich region (Gly, 20–70%) at the C-terminus [31]. The GRPs harboring characteristic internal CCHC-type zinc finger are designated as RZ-1 proteins and include three members in *Arabidopsis thaliana* (AtRZ-1A, AtRZ-1B, and AtRZ-1C) [32–34]. Evidence is increasing that GRPs are linked to plant growth and development as well as stress responses by regulating transcriptional and post-transcriptional processes [31]. For example, *A. thaliana* RZ-1B and RZ-1C interact with a spectrum of splicing-related proteins and directly bind to FLOWERING LOCUS C (FLC) chromatin to regulate splicing of *FLC* introns [35]. Of the GRPs, GRP7 is the most intensively studied in *A. thaliana*, and it controls the transcription level of oscillation-related genes in a clock-regulated negative feedback circuit [36]. In addition, *A. thaliana* GRP8, the paralog of GRP7, can regulate AS events via direct interaction with transcripts [37]. Although detailed mechanism remains unknown, current findings provide sufficient evidence that GRPs participate in transcriptional and post-transcriptional regulation in cold, salt, and dehydration stress [33,38,39].

Based on advances in high-throughput transcriptome sequencing, up to 70% of all plant multi-exon genes are estimated to undergo AS and most AS events require the spliceosome [40]. The spliceosome consists of five snRNAs (U1, U2, U4, U5, and U6) and nearly 300 proteins. U1-70K directly interacts with U1 snRNA to recognize the conserved GU dinucleotide at the 5′ donor site of intron and act as a signal for accurate splicing of pre-mRNAs [41, 42]. Combined effect of GRP7, GRP8, U1-70K, and other spliceosomal proteins constitutively regulate pre-mRNA AS to alter the transcriptome [41]. Although plant spliceosomes have not been isolated to date, analysis of the known spliceosomal proteins indicates that core spliceosome machinery is conserved among different species.

In this study, the FYVE domain-containing proteins were systematically identified and analyzed in oomycete, and one secreted FYVE domain-containing protein from *P. sojae* (PsFYVE1) was selected for further study. PsFYVE1 was essential for *P. sojae* virulence and suppressed plant immunity. PsFYVE1 bound to PI3P that is enriched during *Phytophthora* infection via its conserved FYVE domain. PsFYVE1 targeted RZ-1A in *N. benthamiana* and soybean, and silencing of *NbRZ-1A* increased *Phytophthora* infection and suppressed plant defense. GmRZ-1A/NbRZ-1A complementation in *NbRZ-1A*-silenced plants recovered the immunity phenotype. Notably, NbRZ-1A was associated with three key splicing factors, NbGRP7, NbGRP8, and NbU1-70K. PsFYVE1 could disrupt association between NbRZ-1A and NbGRP7. Moreover, according to combined RNA-seq and experimental analyses, PsFYVE1 and NbRZ-1A co-regulated transcription and pre-mRNA AS of immunity-related genes. Overall, the results shed light on the novel *P. sojae* effector PsFYVE1 which can regulate transcription and AS of immunity-related genes by targeting plant RZ-1A proteins to promote *Phytophthora* infection.

## Results

### Expanded FYVE domain-containing protein family exhibits distinct characteristics in oomycetes

The FYVE domain-containing proteins are found in *Homo sapiens* [16], *Saccharomyces cerevisiae* [43], and *A. thaliana* [44], and therefore investigating their occurrence in oomycetes is also intriguing. According to bioinformatics analysis, an average of 109 FYVE proteins was predicted in the genus *Phytophthora*, compared with 99 in *Pythium ultimum*, 88 in *Hyaloperonospora arabidopsidis*, and 97 in *Albugo laibachii* (**Fig 1A**). Among the examined oomycetes, the fish pathogen *S. parasitica* contained the most members with 177. By contrast, none of the five bacterial genomes examined contained FYVE proteins, and much smaller numbers of FYVE proteins were predicted in fungi, plants, and metazoans (**Fig 1A**). Notably, the number of FYVE proteins was not directly correlated with the number of total proteins for a given organism in eukaryotes (**Fig 1A**). Although two diatoms, *Phaeodactylum tricornutum* and *Thalassiosira pseudonana*, were closely related to oomycetes, they contained only four and five FYVE proteins, respectively. Therefore, these suggest that FYVE proteins expanded widely in oomycetes, with some may contribute to pathogen lifestyles (**Fig 1A**).

**Fig 1.**
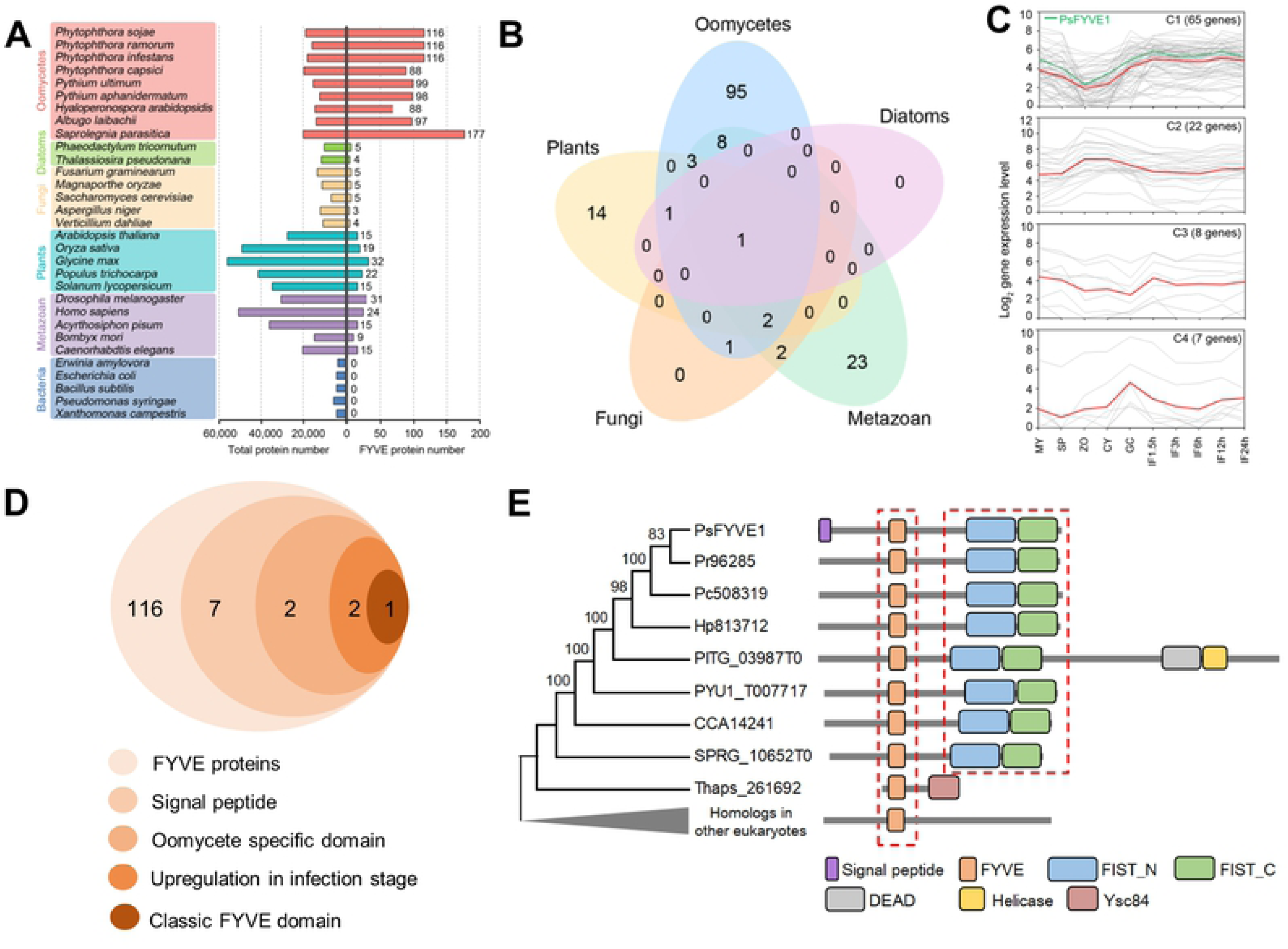
FYVE domain-containing protein family exhibits distinct characteristics in oomycetes. **(A)** Comparison of FYVE domain-containing protein numbers and total protein numbers among representative oomycetes, diatoms, fungi, plants, metazoan, and bacteria. **(B)** Distribution of additional domains in FYVE domain-containing proteins among eukaryotes. **(C)** Four expression patterns of FYVE genes based on *P. sojae* transcriptome data at 10 different developmental and infection stages. **(D)** Flow chart of how to select PsFYVE1 among *P.soaje* FYVE domain-containing proteins. **(E)** Phylogenetic analysis and domain distribution of PsFYVE1 and its orthologs among oomycetes.

To infer the evolutionary history of oomycete FYVE proteins, *P. sojae* was selected to represent the oomycetes, and a phylogenetic analysis of FYVE proteins was conducted with representatives of other eukaryotes. In the phylogenetic tree, only a few *P. sojae* FYVE proteins clustered with homologs from other eukaryotes. Most *P. sojae* FYVE proteins formed independent clades that did not clearly cluster with those of other eukaryotes (**S1 Fig**). The results indicate that some FYVE proteins evolve specifically in oomycetes.

Conservation of FYVE domains in oomycetes was investigated. Notably, 92.5% of oomycete FYVE proteins contained only one FYVE domain, whereas the others contained two to five FYVE domains. According to further sequence analysis, 16.4% of oomycete FYVE domains were of the classic FYVE domain type, but most had variable amino acid composition in the three regions. In addition to the FYVE domain, half of oomycete FYVE proteins had at least one other additional domain. In total, 111 diverse types of domains were identified in oomycete FYVE proteins, significantly more than in proteins of diatoms (2), fungi (6), plants (21), and metazoans (39). Of these domains, oomycetes shared 16 domains with metazoans, seven with plants, four with fungi, and two with diatoms (**Fig 1B**). Notably, 95 domains were found only in oomycetes, and 15 of those were distributed in at least eight of the nine studied oomycetes. These results collectively indicate that oomycete FYVE proteins evolve to exhibit variable FYVE domains and diverse domain architectures. Digital gene expression profiling analysis of *P. sojae* at 10 different developmental and infection stages showed that *P. sojae FYVE* genes were classified into four major clusters (C1 to C4) (**Fig 1C**). The largest cluster was C1, and members had high expression levels at each infection stage compared with those in developmental stages. By contrast, C2 members were highly expressed at zoospore (ZO) and cyst germination (CY) stages. These results suggest that *P. sojae FYVE* genes may have diverse roles during the lifecycle.

To further investigate biological functions of FYVE proteins during interaction between oomycetes and plants, the *P. sojae* FYVE protein PsFYVE1 (Ps136834) was selected based on several criteria and performed subsequent experimental analysis (**Fig 1D**). PsFYVE1 encoded a FYVE protein that contained an N-terminal signal peptide, one classic FYVE domain, and two oomycete-specific FIST domains (**Figs 1E and S2A**). Both RNA-seq and quantitative reverse transcriptase polymerase chain reaction (qRT-PCR) data verified that PsFYVE1 was up-regulated in the infection process (**Figs 1C and S2B**). Further analysis showed that PsFYVE1 was highly conserved in other oomycetes (**Fig 1E**), but had little similarity with those in other eukaryotes. Notably, only PsFYVE1 contained the N-terminal signal peptide, and no polymorphism was found among *P. sojae* isolates, suggesting that PsFYVE1 is unique in *P. sojae*.

### PsFYVE1 contributes to *P. sojae* virulence by suppressing plant defense response

A yeast signal sequence trap system was used to functionally validate the predicted signal peptide of PsFYVE1. The Avr1b signal peptide and the pSUC2 empty vector were used as positive and negative controls, respectively [29, 45]. The PsFYVE1 N-terminal signal peptide sequence was cloned into the yeast vector pSUC2, followed by introduction into invertase secretion-defective yeast YTK12. The YTK12 strain carrying the signal peptide of PsFYVE1 induced secretion of yeast invertase and grew on YPRAA medium (with raffinose instead of sucrose), whereas the YTK12 strain and the YTK12 strain carrying pSUC2 did not induce secretion (**S3 Fig**). Invertase activity was further confirmed by reduction of the dye TTC to the insoluble red color of triphenylformazan (**S3 Fig**). The results suggest that PsFYVE1 carries a functional secretory signal peptide.

To investigate the contribution of PsFYVE1 to *P. sojae* virulence, we utilized a gene silencing technology and successfully obtained three independent *PsFYVE1*-silenced transformants. The qRT-PCR analysis showed that transcript levels of the three silenced transformants (T5, T8, and T15) decreased significantly to 31–42% of those of the WT strain. In the non-silenced transformant T9, transcript levels were not affected (**S4A Fig**). Analysis of mycelial colony growth revealed that filamentous growth in the silenced transformants was similar filamentous to that in the WT strain (**S4B and S4C Fig**). The virulence of the transformants was also tested on etiolated soybean seedlings. Virulence of the three *PsFYVE1*-silenced transformants decreased significantly compared with that of the WT strain (**Figs 2A and 2B**). A relative *P. sojae* biomass (ratio of *P. sojae* DNA to soybean DNA) also indicated that the three silenced transformants were less virulent compared with the WT stain (**Fig 2C**). These results indicate that *PsFYVE1* is an essential virulence factor for *P. sojae* infection.

**Fig 2.**
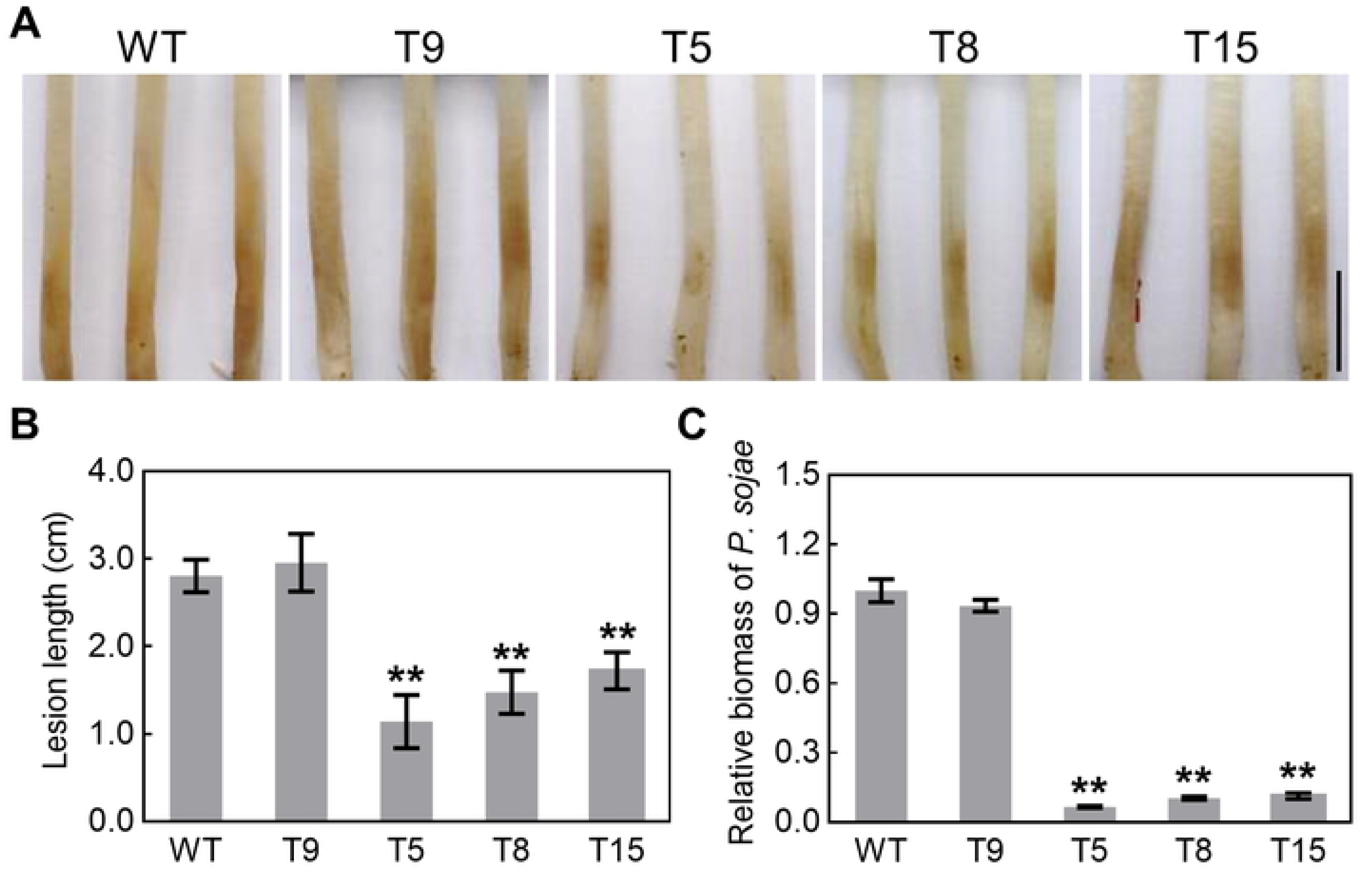
*PsFYVE1*-silenced transformants exhibit significantly reduced virulence. **(A)** Disease symptoms on etiolated hypocotyls infected by *PsFYVE1*-silenced transformants. Wild-type (WT), *PsFYVE1*-silenced transformants (T5/ 8/ 15) and T9 (a non-silenced transgenic line used as a negative control) were incubated on soybean for 36 hpi. Experiments were repeated three times with similar results. Scale bar represents 1 cm. **(B)** Average lesion lengths of the infected etiolated soybean seedlings. Lesion lengths (cm) were measured at 36 hpi. **(C)** Relative biomass quantification of the inoculated soybean seeding. DNA from *P. sojae* infected regions was isolated at 36 hpi and primers specific for the soybean and *P. sojae* actin genes were used for qRT-PCR. All data are shown as mean ± _SD_ (n = 3-6). Asterisks indicate statistical significance: **P<0.01, two-sided unpaired *t* test.

To explore the virulence functions of secreted PsFYVE1 in planta, *N. benthamiana* leaves overexpressing signal peptide-removed PsFYVE1-GFP or GFP alone (negative control) were inoculated with *P. capsici*. Expression of PsFYVE1-GFP in *N. benthamiana* significantly promoted *P. capsici* infection, compared with the GFP control (**Figs 3A and 3C**). The stability of PsFYVE1-GFP and GFP fusion proteins was detected by western blotting (**Fig 3B**). Consistently, the relative *P. capsici* biomass with PsFYVE1-GFP overexpression was higher than that in the GFP control (**Fig 3D**). Taken together, these results reveal that PsFYVE1 can impair plant defense and promote *P. capsici* infection. As the activation of plant immune responses is often accompanied by accumulation of reactive oxygen species (ROS), and successful pathogens may deploy virulence factors that conquer such plant basal resistance [46–48]. Here, flg22- and chitin-triggered ROS were selected as examples, and responses to PsFYVE1 were examined. As shown in **Fig 3E**, flg22-induced ROS production in PsFYVE1-expressing leaves was reduced by 74% compared with that in leaves with the empty vector. Similarly, chitin treatment generated a 63% reduction in ROS accumulation in PsFYVE1-overexpressing leaves compared with that in leaves with the empty vector (**Fig 3F**). Thus, these results suggest that PsFYVE1 can suppress plant defense by inhibiting ROS accumulation.

**Fig 3.**
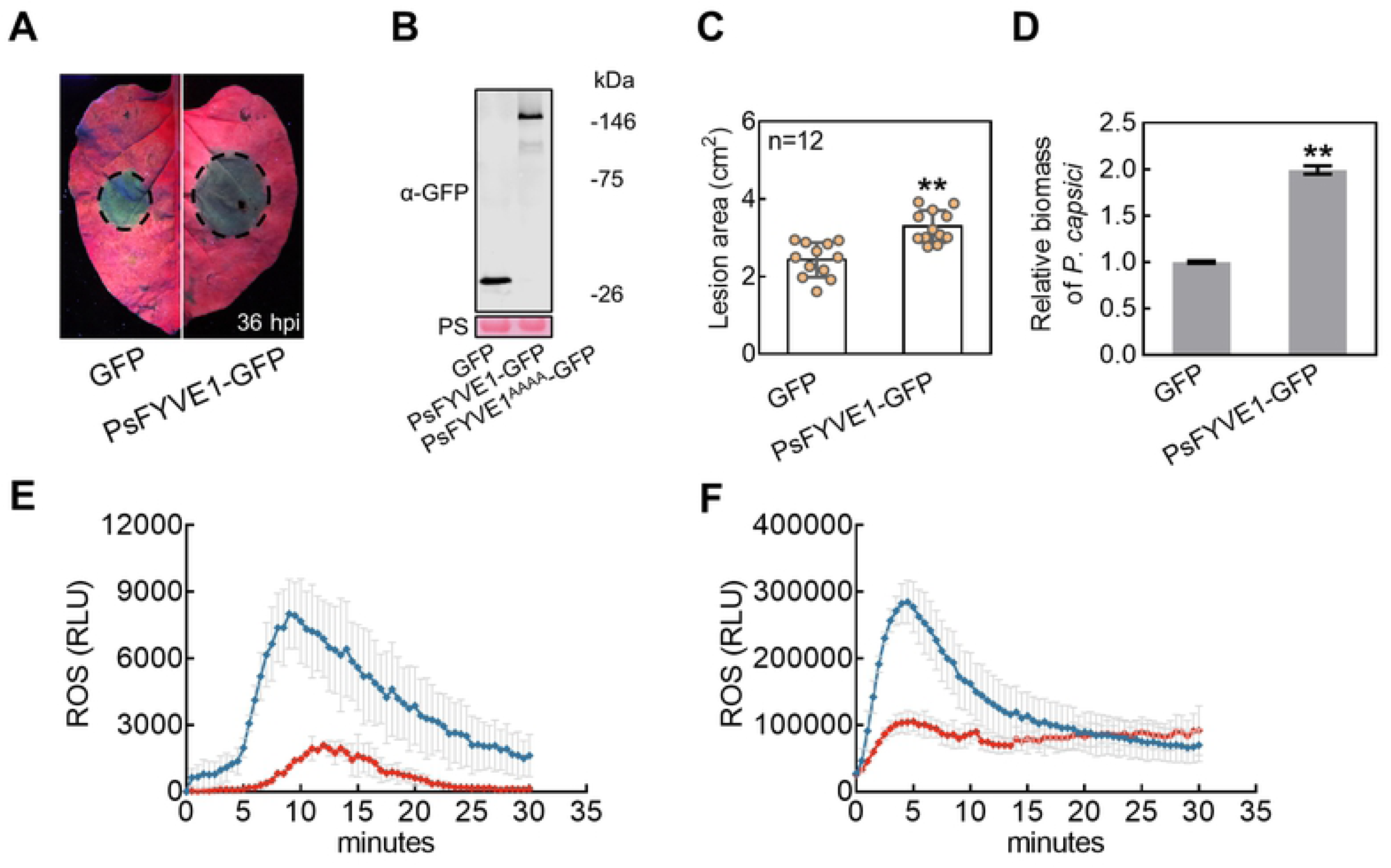
PsFYVE1 promotes *Phytophthora* infection and inhibits plant immunity. **(A)** Conserved residues of the FYVE motif are important for PsFYVE1 induced susceptibility. The *N. benthamiana* leaves were infiltrated to express the indicated proteins, and then inoculated with *P. capsici* 24 hours later. Photographs were taken at 36 hpi under UV light. Dashed lines indicate the lesion areas. **(B)** Protein expression of validated by western blotting using an anti-GFP antibody. Protein loading is visualized by Ponceau stain. **(C)** Average lesion sizes of infected *N. benthamiana* leaves. Lesion area (cm^2^) was measured at 36 hpi. Error bars represent the mean ± _SD_ (n = 12). Asterisks indicate statistical significance: **P<0.01, two-sided unpaired *t* test. **(D)** Relative biomass quantification of inoculated *N. benthamiana* leaves. DNA from *P. capsici* infected regions was isolated at 36 hpi and primers specific for the *N. benthamiana* and *P. capsici* actin genes were used for qRT-PCR. qRT-PCR was performed and normalized to GFP (**P<0.01, two-sided unpaired *t* test). **(E-F)** Flg22- and chitin-induced reactive oxygen species (ROS) burst in the GFP and PsFYVE1-GFP overexpressed *N. benthamiana* leaves. Relative luminescence units (RLU) indicate relative amounts of H_2_O_2_ production induced by 1 μM flg22 (E) and 100 μg/mL chitin (F) in leaf discs. The results are representative of three independent experiments. Error bars represent the mean ± _SD_ (n = 9).

### FYVE domain of PsFYVE1 has PI3P-binding activity

As the FYVE zinc finger domain usually functions in binding to PI3P [49, 50], whether PsFYVE1 and its FYVE domain could bind to PI3P and other phosphoinositides was tested. The R and H residues in the FYVE domain provide critical hydrogen bonds to PI3P [16, 22]. Therefore, a *PsFYVE1^AAAA^* mutant gene was synthesized with the first R and two H residues of the R(R/K)HHCR motif and the R of the R(V/I)C motif substituted into A (**S5A Fig**). In addition, a construct encoding a tandem repeat of the PsFYVE1 FYVE domain (2×FYVE) was generated to test phosphoinositide-binding activities. Six PIs were spotted onto Hybond-C membranes, which were incubated with purified protein of GST fused to the N-terminal full-length PsFYVE1, PsFYVE1^AAAA^, or 2×FYVE. The mouse FYVE protein, EEA1, was the positive control; whereas the empty vector was the negative control. As shown in **S5B Fig**, GST-PsFYVE1 and GST-2×FYVE bound to PI3P, whereas GST-PsFYVE1^AAAA^ did not bind, suggesting that PsFYVE1 specifically bind to PI3P on the basis of the conserved amino acids of the FYVE domain. Furthermore, GST-PsFYVE1, GST-PsFYVE1^AAAA^, and GST-2×FYVE could also bind to PI5P, whereas GST-EEA1 could not (**S5B Fig**).

### PsFYVE1 interacts with plant RZ-1A, RZ-1B, and RZ-1C through its FIST_C domain

To identify plant targets of PsFYVE1, PsFYVE1-GFP was transiently expressed in *N. benthamiana* leaves followed by immunoprecipitation (IP) assay using GFP-Trap_A beads with GFP as the control. Immuno-purified proteins were then analyzed by LC-MS/MS, and 52 *N. benthamiana* proteins in parallel repetitions potentially interacted with PsFYVE1 (**S1 Table**). Based on sequence similarity, the largest group of proteins extensively linked with the DNA-/RNA-binding processes and were selected for further analysis. Of the DNA-/RNA-binding proteins, the homolog of NbRZ-1B in *A. thaliana* participates in plant growth and development [35], but its role in plant immunity remains unknown. Therefore, NbRZ-1B was selected as the potential target for subsequent study.

NbRZ-1B is in the zinc finger-containing glycine-rich RNA-binding protein family, which has three members in *A. thaliana* (AtRZ-1A, AtRZ-1B, and AtRZ-1C) [32,35,51]. Bioinformatics analysis showed that NbRZ-1A and NbRZ-1C were the paralogs of NbRZ-1B. To confirm the interactions between PsFYVE1 and NbRZ-1A/1B/1C, a Co-IP assay was performed. The results clearly showed that PsFYVE1 was significantly enriched in NbRZ-1A/1B/1C precipitates but not in that of the GFP control (**S6A Fig**). The interactions were further validated via the bimolecular fluorescence complementation (BiFC) experiments. NbRZ-1A/1B/1C (NbRZ-1A-Yn, NbRZ-1B-Yn and NbRZ-1c-Yn) and PsFYVE1 (PsFYVE1-Yc) were co-expressed in *N. benthamiana*. YFP fluorescence was observed exclusively in the nuclear speckles, whereas YFP fluorescence was not detected in the negative control (**S6B Fig**). Taken together, these experiments demonstrate that PsFYVE1 interacts with NbRZ-1A/1B/1C and that the proteins co-localize in nuclear speckles.

To test whether PsFYVE1 also interacted with NbRZ-1A/1B/1C homologs in soybean, the natural host of *P. sojae*, we cloned *GmRZ-1A*/*1B*/*1C* full-length cDNA sequences using *Glycine max*, cv. William 82 cultivar. The interactions between GmRZ-1A/1B/1C and PsFYVE1 were validated via the Co-IP experiment (**Fig 4A**) and the BiFC assay (**Fig 4B**). Moreover, the interactions were tested using a luciferase complementation assay. When PsFYVE1 was fused with nLUC at the C-terminal, GmRZ-1A/1B/1C fused with cLUC at the N-terminal showed clear interactions with cLUC-PsFYVE1 in *N. benthamiana* leaves (**Fig 4C**). No luminescent signal could be detected in negative controls (**Fig 4C**). Therefore, all evidence demonstrates that RZ-1A/1B/1C proteins in *N. benthamiana* and soybean are the targets of PsFYVE1.

**Fig 4.**
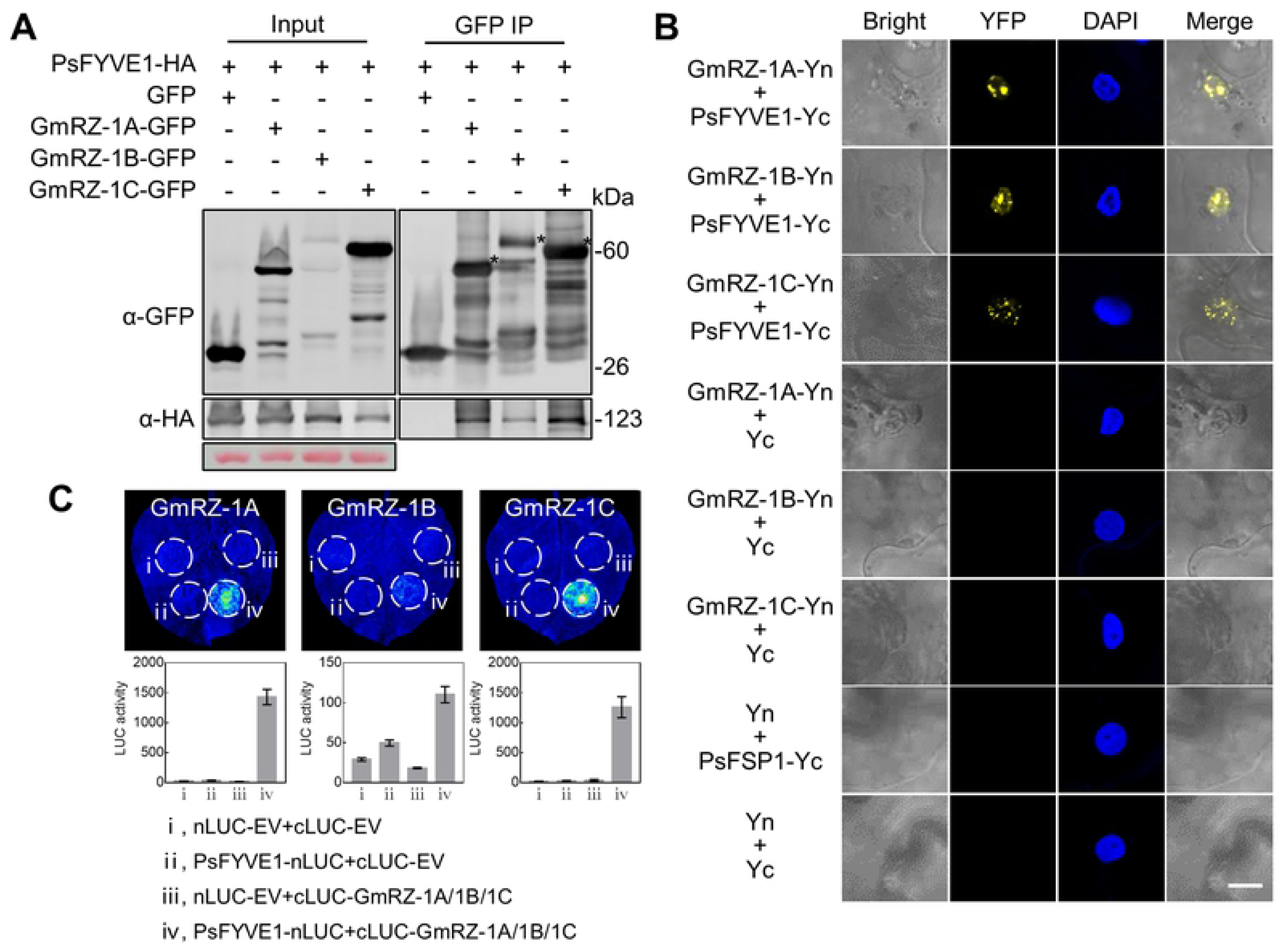
PsFYVE1 interactes with GmRZ-1A, GmRZ-1B and GmRZ-1C in vivo. **(A)** Co-immunoprecipitation (IP) assay of PsFYVE1 and GmRZ-1A/1B/1C. IP of protein extracts from agroinfiltrated leaves using GFP-Trap confirmed that PsFYVE1-HA was associated with GmRZ-1A/1B/1C, but not the GFP control. The positions of the expected protein bands are indicated by asterisks. Protein sizes are indicated in kilodaltons (kDa), and protein loading is visualized by Ponceau stain. **(B)** Interaction and co-localization of PsFYVE1 and GmRZ-1A/1B/1C in bimolecular fluorescence complementation (BiFC) assay. *A. tumefaciens* cells harboring the indicated YFPn (Yn)- or YFPc (Yc)-fused constructs were co-infiltrated into *N. benthamiana* leaves. The fluorescent signals were detected by confocal laser microscopy at 48 hours after infiltration and 4’6-diamidino-2-phenylindole (DAPI) were used to a nucleus staining. Scale bar represents 20 μm. **(C)** Association of PsFYVE1 and GmRZ-1A/1B/1C in split-luciferase complementation assay. Leaves were used to measure the LUC activity and twenty-four leaf discs were used to measure the luminescence 48 h after co-expression of the indicated proteins. Error bars represent the mean ± _SD_ (n = 24).

To determine the precise subsections of PsFYVE1 that dominated its interactions with RZ-1 proteins, four PsFYVE1 residue-exchange and deletion mutants were constructed (**Fig 5A**). In the luciferase complementation assay, the luminescence signal was not detected in leaves expressing PsFYVE1^ΔFIST_N^-nLUC or PsFYVE1^ΔFIST_C^-nLUC with cLUC-GmRZ-1A/cLUC-NbRZ-1A (**Fig 5A**). LUC activity measurements also confirmed those results (**Fig 5B**). Thus, the FIST_N and FIST_C domains, rather than the FYVE domain, are essential for interactions between PsFYVE1 and RZ-1A proteins.

**Fig 5.**
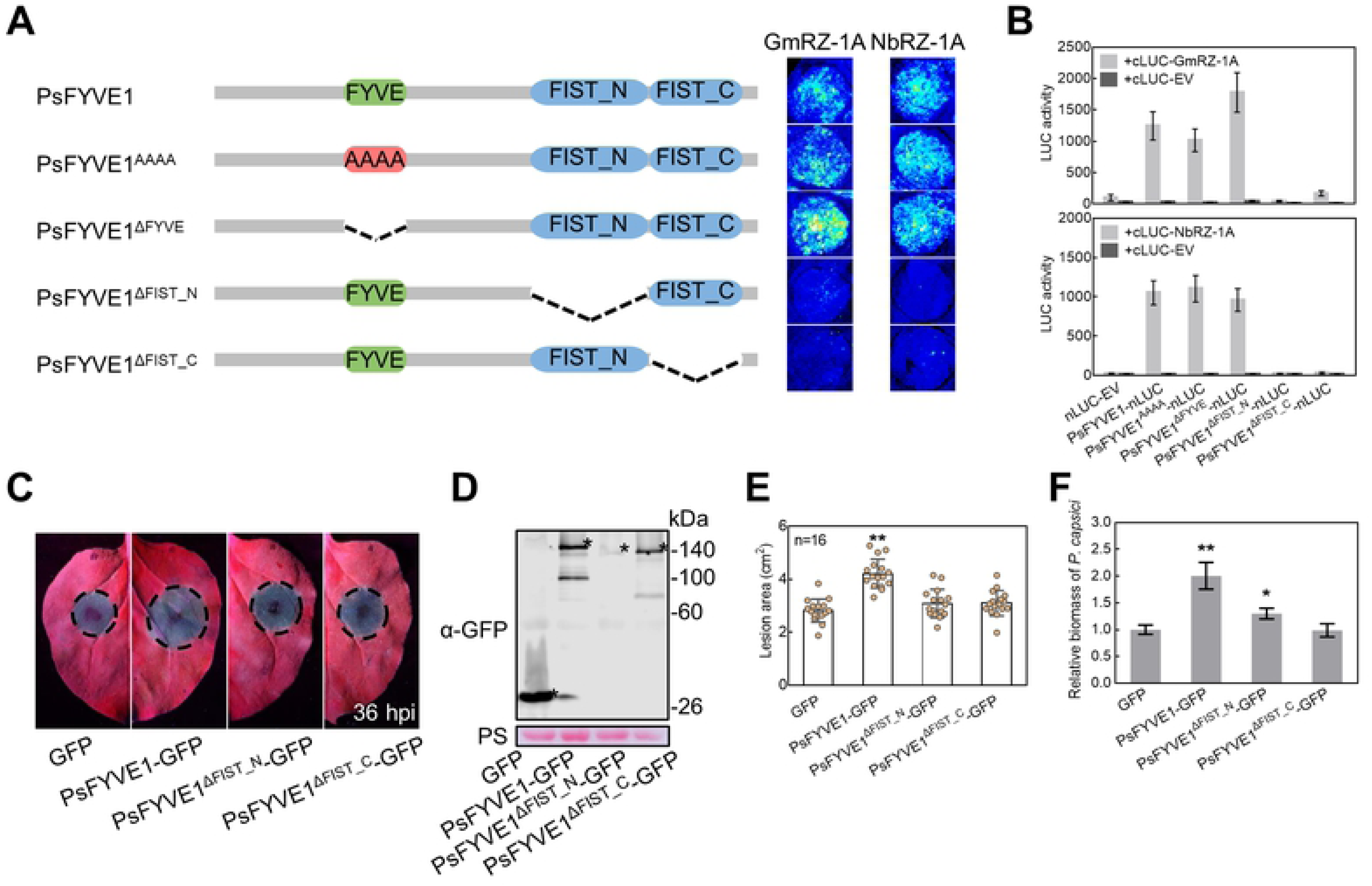
FIST domain is essential for the interaction between PsFYVE1 and RZ-1A as well as the promotion of *Phytophthora* infection. (A) Verifying the interaction between PsFYVE1 mutants and RZ-1A in split-LUC assays. Schematic drawings of PsFYVE1 and mutants are shown on the left. The interaction between PsFYVE1 mutants and RZ-1A is shown on the right. **(B)** Twelve leaf discs were used to measure the luminescence 48 h after co-expression of the indicated proteins. Error bars represent the mean ± _SD_ (n = 12). (C) *P. capsici* infection assay on leaves expressing PsFYVE1-GFP, PsFYVE1^ΔFIST_N^-GFP or PsFYVE1^ΔFIST_C^-GFP. *P. capsici* mycelia were inoculated on the leaves at 24 hours after Agro-infiltration. Photographs were taken at 36 hpi under UV light. Dashed lines indicate the lesion areas. **(D)** Protein expression of GFP, PsFYVE1-GFP PsFYVE1^ΔFIST_N^-GFP and PsFYVE1^ΔFIST_C^-GFP in western blotting. The positions of the expected protein bands are indicated by asterisks. Molecular weight markers are indicated in kDa, and protein loading is visualized by Ponceau stain. **(E)** Average lesion sizes of infected *N. benthamiana* leaves. Lesion sizes (cm^2^) were measured at 36 hpi. Data are shown as mean ± _SD_ (n = 16). Asterisks indicate statistical significance: **P<0.01, two-sided unpaired *t* test. **(F)** Relative biomass quantification of inoculated *N. benthamiana* leaves. DNA from *P. capsici* infected regions was isolated at 36 hpi and primers specific for the *N. benthamiana* and *P. capsici* actin gene were used for qRT-PCR. qRT-PCR was performed and normalized to GFP (** P<0.01, * P<0.05; two-sided unpaired *t* test).

We found that ectopic expression of the PsFYVE1^ΔFIST_N^ mutant in *N. benthamiana* did not enhance plant susceptibility to *P. capsici*, and its protein accumulation level was decreased significantly (**Figs 5C and 5D**). Moreover, the FIST_C deletion mutant did not enhance *N. benthamiana* susceptibility, and protein expression was stable (**Figs 5C and 5D**). The lesion areas and relative biomass assay also confirmed the results (**Figs 5E and 5F**). Therefore, PsFYVE1 promotes *Phytophthora* infection via its linkage to GmRZ-1A/NbRZ-1A, which depends on its FIST_C domain.

### Disease resistance and ROS are attenuated in NbRZ-1A-silenced leaves, which can be complemented by GmRZ-1A and NbRZ-1A

*A. thaliana* RZ-1 proteins are known to participate in abiotic stress responses [51, 52], ABA signaling [53] and plant development [35]. However, whether RZ-1 proteins are associated with plant defenses is largely unknown. To further characterize the roles of RZ-1 proteins in defense response, *NbRZ-1A*, *NbRZ-1B*, and *NbRZ-1C* genes in *N. benthamiana* were silenced using *Agrobacterium*-mediated VIGS technology. Silencing efficiency of *NbRZ-1A*, *NbRZ-1B*, and *NbRZ-1C* ranged from 64 to 51% that was indicated by qRT-PCR data (**Fig 6A**). However, TRV:NbRZ-1C was not specifically silenced and interfered with the *NbRZ-1B* gene (**Fig 6A**). Silencing of NbRZ-1B and NbRZ-1C resulted in stunted plants with smaller leaves, whereas silencing of NbRZ-1A did not alter growth compared with that in TRV:GUS (negative control) plants (**Fig 6B**). Then *P. capsici* infection assay was conducted on silenced leaves. *NbRZ-1A*/*1B*/*1C*-silenced plants showed significantly larger disease lesions than *GUS*-silenced control (**Figs 6B and 6C**), and relative biomass assay also confirmed those results (**Fig 6D**). Afterward, effects of silencing *NbRZ-1A*/*1B*/*1C* on ROS accumulation triggered by flg22 and chitin was tested. As shown in **Fig 6E**, flg22-induced ROS production in *NbRZ-1A*/*1B*/*1C*-silenced leaves decreased by 67%, 66%, and 80%, respectively, compared with leaves in the *GUS*-silenced control. Similarly, chitin treatment generated 79%, 74%, and 54% reduction in ROS accumulation in *NbRZ-1A*/*1B*/*1C*-silenced leaves compared with leaves in the *GUS*-silenced control (**Fig 6F**). Thus, the results suggest that *NbRZ-1A*/*1B*/*1C* are positive regulators of plant immunity.

**Fig 6.**
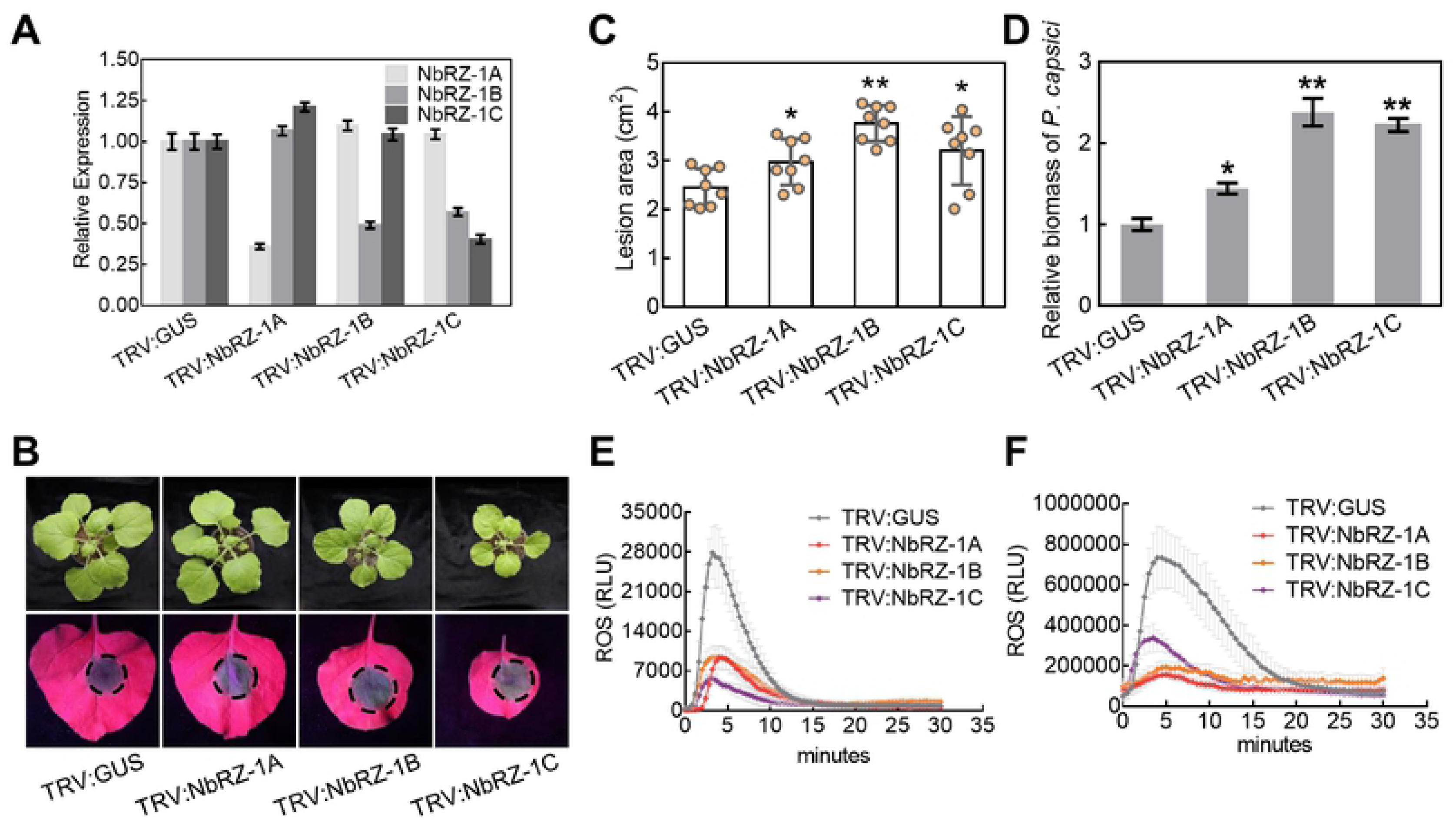
Silencing of *NbRZ-1* in *N. benthamiana* attenuates plant immunity. **(A)** Lesions caused by *P. capsici* on the *NbRZ-1A*/*1B*/*1C*-silenced *N. benthamiana* leaves. The silenced-plants were photographed at 30 days after infiltration (upper panel). *P. capsici* infection photographed on leaves (down panel) were taken at 36 hpi under UV light. The dashed lines indicate the lesion areas. The experiments were replicated three times with similar results. **(B)** Silencing efficiency of *NbRZ-1A*/*1B*/*1C* genes. The qRT-PCR results show that each of three genes is significantly silenced by virus-induced gene silencing (VIGS) technology. TRV:GUS was used as a control. Data are shown as means ± _SD_ (n = 3). **(C)** The average lesion areas caused by *P. capsici* at 36 hpi. The data are shown as mean ± _SD_ (n = 8). Asterisks indicate statistical significance: **P<0.01; * P<0.05, two-sided unpaired *t* test. **(D)** Relative *P. capsici* biomass quantified by qPCR. DNA from infected regions was isolated at 36 hpi for qPCR analyses. Means and standard deviations from three replicates were shown (** P<0.01; * P<0.05, two-sided unpaired *t* test). **(E-F)** Suppression of flg22- and chitin-induced ROS production in the NbRZ-1*A*/*1B*/*1C*-silenced leaves. Relative luminescence units (RLU) indicate the relative amounts of H_2_O_2_ production induced by 1 μM flg22 (E) and 100 μg/mL chitin (F). Means and standard deviations from three replicates are shown.

Based on public RNA-seq data [54, 55], transcription level of *GmRZ-1A* and *NbRZ-1A* are higher than those of *GmRZ-1B*/*1C* and *NbRZ-1B*/*1C* (**S7A Fig**). Therefore, *GmRZ-1A* and *NbRZ-1A* were investigated further in this study. According to multiple sequence alignments of full-length amino acid sequences of GmRZ-1A and NbRZ-1A, GmRZ-1A shared 64% identity with NbRZ-1A, and both contained a glycine, arginine, and aspartic acids-rich region in the C-terminus (**S7B Fig**) consistent with *A. thaliana* RZ-1 proteins [35, 56].

To further confirm silencing NbRZ-1A caused the susceptible phenotype, the synthetic GmRZ-1A^syn^ or NbRZ-1A^syn^, synonymous mutation of GmRZ-1A or NbRZ-1A that was transiently expressed to complement the *NbRZ-1A*-silenced *N. benthamiana* leaves. After *P. capsici* challenge, overexpression of GmRZ-1A^syn^-GFP and NbRZ-1A^syn^-GFP suppressed *NbRZ-1A* silencing-mediated infection promotion and ROS inhibition (**S8A and S8D Figs**). Leaves with GmRZ-1A^syn^/NbRZ-1A^syn^ developed significantly smaller lesion than those in leaves with the GFP control in *NbRZ-1A*-silenced *N. benthamiana* (**S8B Fig**). Western blotting was used to analyze the protein stability (**S8C Fig**). The results suggest that NbRZ-1A and GmRZ-1A act as orthologous proteins and maintain an identical positive regulatory role in plant immunity.

### RZ-1A C-terminus is essential for interaction with PsFYVE1 and immune function

To test the relation between immune-regulated function and different domains of RZ-1A, a series of truncated versions of RZ-1A were generated, including GmRZ-1A^ΔCT^ (amino acids 1–110), GmRZ-1A^ΔRRM^ (amino acids 80–208), NbRZ-1A^ΔCT^ (amino acids 1–110), and NbRZ-1A^ΔRRM^ (amino acids 78–209) (**S9A Fig**). To determine the precise subsections of RZ-1A that dominated the interaction of PsFYVE1 and GmRZ-1A/NbRZ-1A, luciferase complementation assays were performed. The results showed that GmRZ-1A^ΔRRM^ and NbRZ-1A^ΔRRM^ interacted with PsFYVE1 but GmRZ-1A^ΔCT^ and NbRZ-1A^ΔCT^ did not (**S9B Fig**), suggesting the interaction of RZ-1A and PsFYVE1 depends on the C-terminus of RZ-1A proteins. Then, *P. capsici* was inoculated on the *N. benthamiana* leaves transiently expressing the above RZ-1A mutants. GmRZ-1A^ΔCT^ and NbRZ-1A^ΔCT^ did not reduce *P. capsici* infection compared with GmRZ-1A/NbRZ-1A (**S9C Fig**). Lesion areas with GmRZ-1A^ΔCT^ and NbRZ-1A^ΔCT^ overexpression were not statistically different from those of the GFP control (**S9D Fig**). Stability and size of each fusion protein were checked by western blotting (**S9E Fig**). In addition, that GmRZ-1A^ΔCT^ and NbRZ-1A^ΔCT^ mutants altered nuclear speckles localization of GmRZ-1A-GFP and NbRZ-1A-GFP (**S9F Fig**). The results confirm that the RZ-1A C-terminus is essential not only for its interaction with PsFYVE1 but also for its function in immune regulation.

### PsFYVE1 disrupts association of NbRZ-1A and splicing factors

Despite potential roles of *A. thaliana* RZ-1B and RZ-1C in pre-mRNA splicing and general gene expression, reports demonstrating transcription and splicing functional roles of RZ-1A proteins are very limited [35, 57]. Therefore, we expressed NbRZ-1A in *N. benthamiana* and its binding proteins was harvested by co-immunoprecipitation. Immunoprecipitation samples were in-gel separated and subjected to LC-MS/MS analyses. A total of 51 *N. benthamiana* proteins in parallel repetitions were identified to potentially interact with NbRZ-1A (**S2 Table**). Among them, peptides matching glycine-rich RNA binding protein 7 (NbGRP7) was the ortholog of *A. thaliana* GRP7, which is a well-studied plant spliceosomal protein.

In previous studies, *A. thaliana* GRP7 associates with its homolog GRP8 or core spliceosome component U1-70K to modulate dynamic AS [28,58–60]. Therefore, full-length cDNAs of *N. benthamiana NbGRP7* (*Niben101Scf03953g01007.1*), *NbGRP8* (*Niben101Scf09268g00007.1*), and *NbU1-70K* (*Niben101Scf05610g01007.1*) were cloned, and their interactions with NbRZ-1A were investigated by luciferase complementation (**Fig 7A**), Co-IP (**Fig 7B**), and BiFC (**Fig 7C**) assays. The results showed that NbRZ-1A interacted with components of the spliceosome complex and likely participated in gene transcriptional and post-transcriptional regulations.

**Fig 7.**
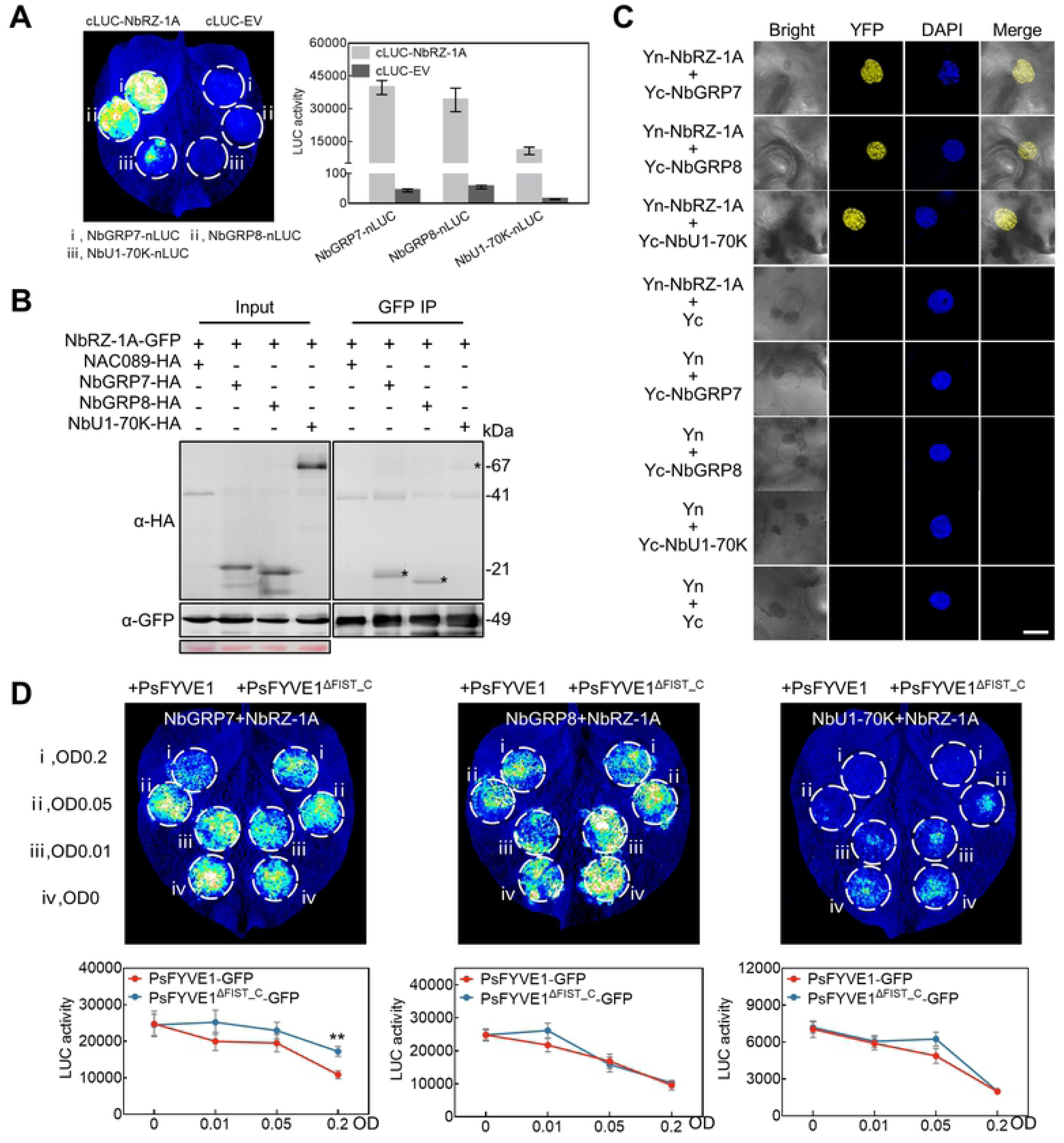
PsFYVE1 disrupts association of NbRZ-1A and plant splicing factor. **(A)** Association of NbRZ-1A with *N. benthamiana* GRP7, GRP8, and U1-70K in split-LUC assay. Leaves and fifteen leaf discs were used to measure the LUC activity and 48 h after co-expression of the indicated proteins. Error bars represent the mean ± _SD_ (n = 15). **(B)** Co-IP assay of PsFYVE1 and NbGRP7/NbGRP8/NbU1-70K. IP of protein extracts from agroinfiltrated leaves using GFP-Trap confirmed that NbRZ-1A-GFP was associated with HA-tagged NbGRP7, NbGRP8, and NbU1-70K, but not the NAC089-HA (HA-tagged the Arabidopsis nucleus-localized protein) negative control. The positions of the expected protein bands are indicated by asterisks. Protein sizes are indicated in kDa, and protein loading was visualized by Ponceau stain. **(C)** Interaction and co-localization of NbRZ-1A and NbGRP7/NbGRP8/NbU1-70K in BiFC assay. *A. tumefaciens* cells harboring the indicated YFPn (Yn)- or YFPc (Yc)-fused constructs were co-infiltrated into *N. benthamiana* leaves. The fluorescent signals were detected by confocal laser microscopy and DAPI were used to a nucleus staining. Photographs were taken 48 hours after infiltration. Scale bar represents 20 μm. **(D)** Interference of interaction between NbRZ-1A and plant splicing factors by PsFYVE1. cLUC-NbRZ-1A and NbGRP7/NbGRP8/NbU1-70K-nLUC were co-expressed in *N. benthamiana* leaves in the presence of PsFYVE1-GFP or PsFYVE1^ΔFIST_C^-GFP. The concentrations (OD_600_) of Agrobacteria carrying PsFYVE1-GFP or PsFYVE1^ΔFIST_C^-GFP were 0, 0.01, 0.05, and 0.2 respectively. PsFYVE1^ΔFIST_C^-GFP indicates a mutant described in supplemental Figure 13A and is used as a control. All data are shown as mean ± _SD_ (n = 9).

To test PsFYVE1 effects on the spliceosome complex of NbRZ-1A–NbGRP7, NbGRP8, and NbU1-70K complex, luciferase complementation assays of NbRZ-1A and NbGRP7/NbGRP8/NbU1-70K were performed in the presence of different concentrations of PsFYVE1-GFP, with non-interacting PsFYVE1^ΔFIST_C^-GFP as the negative control. We observed that the luminescence signal of cLUC-NbRZ-1A and NbGRP7-nLUC binding gradually decreased with the increase in concentrations of PsFYVE1-GFP. The luminescence signal of cLUC-NbRZ-1A and NbGRP8/NbU1-70K-nLUC binding also decreased as the concentration of PsFYVE1-GFP/PsFYVE1^ΔFIST_C^-GFP increased, but the signal was not significantly different at the same concentration of PsFYVE1-GFP/PsFYVE1^ΔFIST_C^-GFP (**Fig 7D**). Therefore, we conclude that PsFYVE1 can disrupt the association of NbRZ-1A and NbGRP7.

### PsFYVE1 and NbRZ-1A co-regulate transcription of defense-related genes

On the basis of above results, RNA-seq technology was used to further explore whether PsFYVE1 and NbRZ-1A regulate transcription and alternative pre-mRNA splicing *in planta*. *N. benthamiana* leaves overexpressing PsFYVE1 and overexpressing GFP (control) and *NbRZ-1A*-silenced and *GUS*-silenced (control) leaves were inoculated with *P. capsici* for 36 hpi, followed by subsequent RNA sequencing. A total of 79 differentially expressed genes (DEGs) were identified in both PsFYVE1-overexpressing and *NbRZ-1A*-silenced samples (**S10A Fig and S3 Table**). Among them, 60 genes were up-regulated and 19 genes were down-regulated. GO enrichment analysis indicated that the 79 DEGs primarily participated in immune system process, response to stimulus, metabolic process, and regulation of biological process (**S10B Fig**).

Among these, three defense-related DEGs were selected: a transferase of plant secondary metabolites (*NbHCT*), a central factor in ET pathways (*NbEIN2*), and a sucrose-cleaving enzyme (*NbSUS4*) for validation of transcriptional changes by qRT-PCR analysis. The qRT-PCR results showed that PsFYVE1 overexpression or *NbRZ-1A* silencing promoted *NbHCT* transcription but repressed *NbEIN2* and *NbSUS4* transcription (**S10C, S10D and S10E Figs**), which were results consistent with RNA-seq data. In previous studies, it has been proven that HCT can subvert the expression of pathogenesis-related (PR) genes by modifying lignin content and composition [61], and therefore may be a negative regulator of plant immunity. By contrast, EIN2 and SUS4 are positive regulators of plant immunity. *A. thaliana* EIN2 is involves in trafficking signal inducing-ethylene response, and its deficient mutant enhances susceptibility to *Macrophomina phaseolina* [62]. Another *SUS4* gene can activate plant immune responses against pathogens by modifying sugar metabolism and content [63, 64]. To analyze functions of the three putative defense-related genes, we inoculated *P. capsici* to observe infection in *NbHCT*-, *NbEIN2*-, and *NbSUS4*-silenced *N. benthamiana* leaves. *NbHCT*-silenced leaves were more resistant to *P. capsici*; whereas *NbEIN2*- and *NbSUS4*-silenced leaves were more susceptible (**S10F Fig**). The results indicate that NbHCT negatively regulates immunity to *Phytophthora* attack, whereas NbEIN2 and NbSUS4 are positive regulators of plant immunity to *Phytophthora* attack. Overall, the results suggest that PsFYVE1 interferes with RZ-1A functions to regulate plant immunity by promoting transcription of susceptibility factors and repressing transcription of positive regulators of plant immunity.

### PsFYVE1 and NbRZ-1A modulate pre-mRNA alternative splicing of defense-related genes

Notably, we found 120 AS events were differentially spliced in the above RNA-seq data, suggesting that PsFYVE1 and NbRZ-1A co-regulate pre-mRNA AS (**S4 Table**). Among the 120 overlapped differential AS events, skipped exon (SE, 40 events) was the most abundant type, followed by retained intron (RI, 31 events), alternative 3’ splice site (A3SS, 31 events), and alternative 5’ splice site (A5SS, 18 events).

Among the 120 AS events, four were randomly tested, including one SE event and three RI events, by performing qRT-PCR and measuring transcript levels of different isoforms (**Figs 8A and S11**). For *Niben101Scf01240g06009* and *Niben101Scf01702g02006*, splicing ratios (transcript level of the spliced isoform divided by the transcript level of the unspliced isoform, isoform1/2) increased during PsFYVE1 overexpression or *NbRZ-1A* silencing (**S11A and S11B Figs**), whereas that of *Niben101Scf03790g01009* (isoform1/2) decreased (**S11C Fig**). *Niben101Scf01008g02005*, a member of membrane attack complex/perforin (macpf) family (*NbNSL1*), is a defense-related gene in plant immunity [65]. *NbNSL1.1* transcript produces a functional protein of membrane attack complex/perforin (macpf) family, whereas transcript of *NbNSL1.2* undergoes SE, which results in the production of truncated proteins that lacks the membrane-attack complex/perforin (MACPF) domain **(Fig 8A**). Splicing ratios (transcript level of the unspliced isoform divided by the transcript level of the spliced isoform) of *NbNSL1* (*NbNSL1.1*/*NbNSL1.2*) increased in leaves with PsFYVE1-overexpressed or *NbRZ-1A*-silenced (**Fig 8A**).

**Fig 8.**
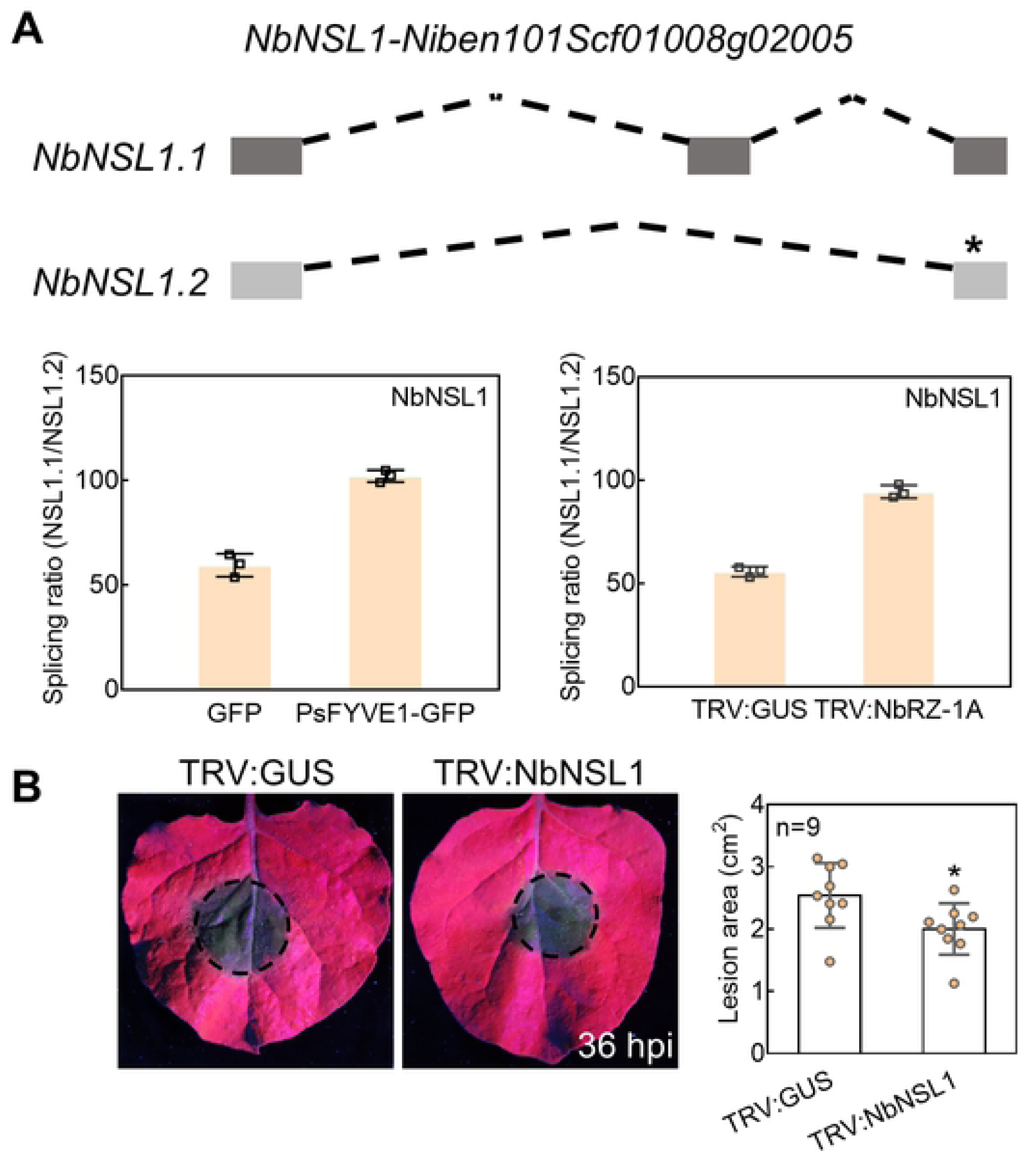
PsFYVE1 and NbRZ-1A are involved in pre-mRNA alternative splicing of the defense-related gene. **(A)** Splicing ratios of the defense-related gene in qRT-PCR analysis. The schematic shows models of two isoforms of *NbNSL1* gene (upper panel), in which the asterisks indicate a premature termination codon. qRT-PCR was performed using isoform-specific primers to measure the splicing ratios of *NbNSL1* gene (lower panel). Total RNA was extracted from the indicated *N. benthamiana* leaf tissues infected with *P. capsici* at 36 hpi, including overexpression of GFP (a control), PsFYVE1-GFP, TRV:GUS (a control), and *NbRZ-1A*-silenced (TRV:NbRZ-1A). Means and standard deviations from three replicates are shown. **(B)** *P. capsici* infection assay on the silenced *N. benthamiana* leaves with silencing of *NbNSL1* gene. *P. capsici* mycelia were inoculated on the leaves at 30 days after agroinfiltration. The photographs and lesion sizes (cm^2^) were taken and measured at 36 hpi. Data are shown as mean ± _SD_ (n = 9). Asterisks indicate statistical significance: * P<0.05, two-sided unpaired *t* test.

NSL1 has been reported to negatively regulate salicylic acid (SA)-mediated pathway of cell death programs and defense responses in *A. thaliana* [65]. To further proceed functional studies of the protein isoforms produced by AS, the putative defense-related genes *NbNSL1* was silenced. *NbNSL1*-silenced leaves were more resistant to *P. capsici* (**Fig 8B**). The results indicated that the NbNSL1 functional isoform was a negative regulator of plant immunity to *Phytophthora* attack. These results all demonstrate that PsFYVE1 interferes with RZ-1A functions to regulate plant immunity by promoting AS of susceptibility factors of plant immunity.

## Discussion

*Phytophthora* pathogens secrete diverse groups of effectors into plants to interfere with plant immune responses and promote infection. Although some types of effectors, such as RXLRs, CRNs, elicitins, and NLPs, have been identified and widely studied, there are still many unknown effectors to be excavated. In this study, an FYVE domain-containing protein family that is highly expanded and with distinct characteristics in oomycetes provided a potential library of novel effectors. The FYVE domain-containing protein PsFYVE1 was selected and functional analysis was performed. We found that PsFYVE1 contained a functional signal peptide and could suppress plant immune response. The conserved FYVE domain of PsFYVE1 is confirmed to be essential for its PI3P-binding activity, suggesting that this class of secreted FYVE proteins could act as novel effectors for *Phytophthora* pathogens. PsFYVE1 interacted with the RNA-binding proteins NbRZ-1A of *N. benthamiana* and GmRZ-1A of host soybean. Silencing of NbRZ-1A in plants suppressed *Phytophthora* infection and ROS accumulation, indicating the NbRZ-1A protein contributes positively to plant resistance. In addition, NbRZ-1A formed a splicing-related complex with GRP7, GRP8, and U1-70K in *N. benthamiana*. PsFYVE1 disrupted association of NbRZ-1A and splicing factors GRP7, suggesting that PsFYVE1–NbRZ-1A interaction interferes with plant immunity at the level of RNA regulation. Both RNA-seq and qRT-PCR analyses indicated that PsFYVE1 and NbRZ-1A co-affected transcription and AS of immune-related genes. Overall, our works reveal that PsFYVE1 is a novel *P. sojae* effector that regulates plant immunity by interfering with the functions of RZ-1A to reprogram gene transcriptional and post-transcriptional process (**Fig 9**).

**Fig 9.**
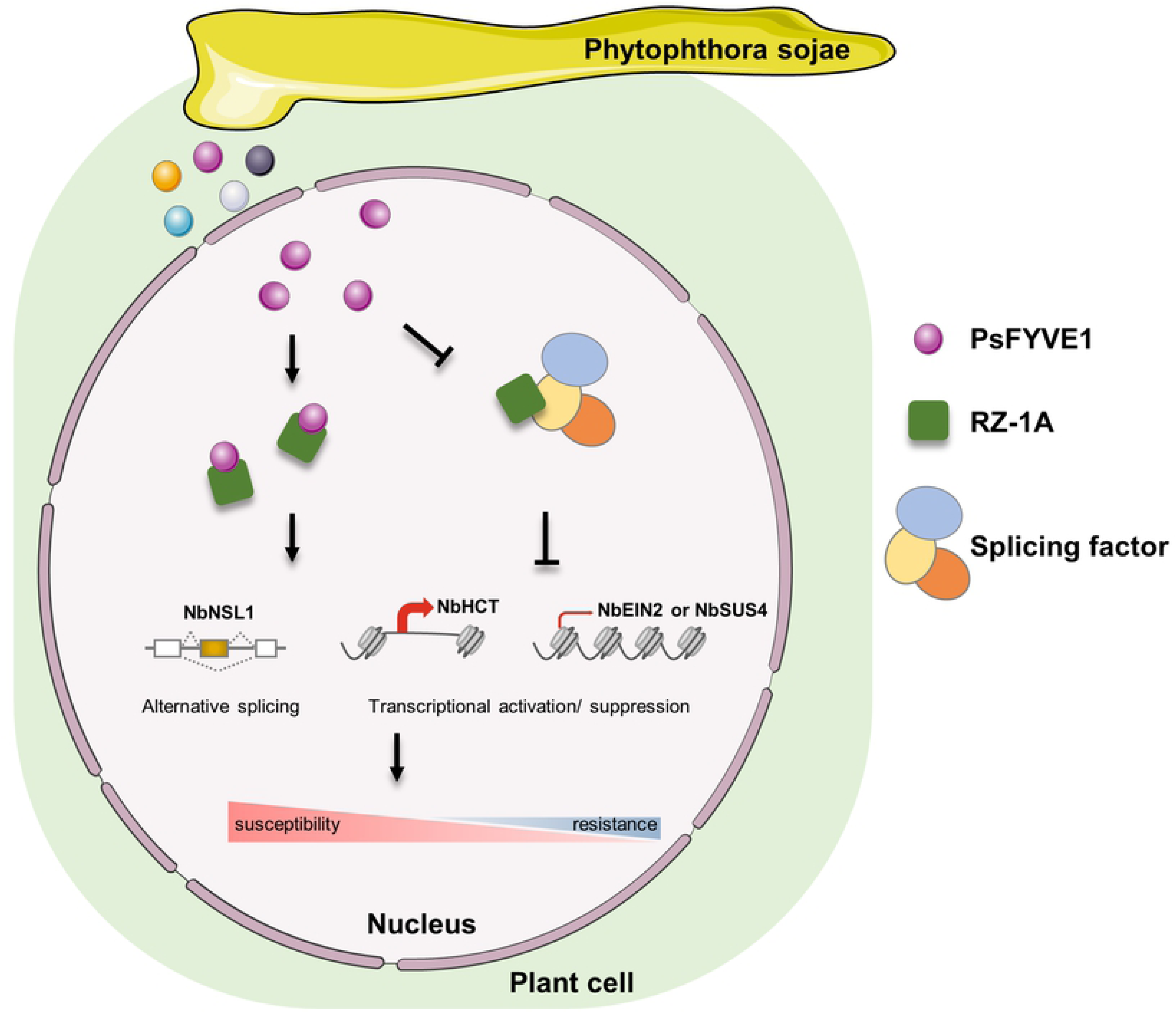
Schematic representation of PsFYVE1 virulence function. A schematic diagram illustrating that PsFYVE1 regulates transcription and alternative splicing through targeting RZ-1A proteins and disrupting RZ-1A-splicing factor interaction. The different colored circles indicate different intracellular effectors and PsFYVE1. One-way solid arrows represent promotion, including interaction of PsFYVE1 and RZ-1A in the presence of PsFYVE1, and inducing transcription (activation of *NbHCT* and suppression of *NbEIN2*/*NbSUS4*) and AS event (exon skipping of *NbNSL1*) to increase plant susceptibility. Solid terminated lines represent inhibition, such as disruption of RZ-1A-splicing factor interaction in the presence of PsFYVE1, and deactivation of transcription and AS. The different colored ovals indicate three splicing factors. The secreted effector PsFYVE1 enter host nucleus, resulting in weak resistance and being more susceptible during infection.

Although the FYVE domain-containing protein family has been widely investigated in humans, yeasts and plants, its distribution, evolution, and functions are poorly known in oomycetes. In this study, FYVE domain-containing proteins were systematically identified and analyzed in oomycetes, especially *P. sojae*. We observed that all the oomycete genomes contained much larger numbers of FYVE proteins compared with other eukaryotes. In addition to conserved FYVE domain, oomycete FYVE proteins also contained a relatively large number of specific additional domains, which might help oomycetes adapt to their distinct environments during evolution. According to published RNA-seq data of *P. sojae*, many FYVE genes were up-regulated at different infection stages, whereas some were highly expressed in developmental stages, suggesting that *P. sojae* FYVE genes may be involved in vegetative growth, stress response, and virulence. In this study, PsFYVE1 contained a signal peptide, and was confirmed to be a novel effector. When the large number of proteins, diverse domain compositions, and variable transcriptional profiles are considered, it is likely that other signal peptide-containing FYVE proteins derived from filamentous pathogens, including oomycetes and fungi, are also potential effectors, which need to be investigated further in the near future.

RXLR effectors secreted from oomycetes can bind with high affinity to PI3P and enter host plant cells via their RXLR motif [13, 20]. As a typical example, *P. sojae* RXLR effector Avr1b is delivered into plant cells through the RXLR and dEER motifs [66]. Whether proteins containing the FYVE domain also have similar functions in membrane transport and effector translocation remains to be determined. Our results showed that PsFYVE1 carried a functional secretory signal peptide and a conserved FYVE domain, which was required for binding to PI3P. The full-length PsFYVE1 and its FYVE domain could bind to PI3P, and this binding depends on four conservative amino acid residues. Thus, we suspect that PsFYVE1 conducts transmembrane transfer to enter into plant cell that depends on the its binding to PI3P. In addition, PsFYVE1 contained FIST_N and FIST_C domains, which are found in signal transduction proteins derived from bacteria, archaea, and eukaryotes [67]. FIST_C domain of PsFYVE1 was essential for its interaction with target proteins, which is consistent with previous study that found FIST domains bind amino acids ligands [67]. Moreover, FIST_C domain of PsFYVE1 is required for PsFYVE1 enhanced susceptibility suggesting that FIST_C specially contributes to virulence function. Overall, PsFYVE1 is a virulence protein secreted by *P. sojae*, and its two functional domains cooperate to perform different functions for successful infection.

RZ-1 proteins including RZ-1A, RZ-1B, and RZ-1C, constitute a small group of plant GRPs that are widespread in higher plants, acting as important regulators of seed maturation, flowering time and freezing tolerance [35,52,53]. However, biological roles of RZ-1 proteins in plant immune response remain unclear. In this study, PsFYVE1 targeted RZ-1A/1B/1C proteins in *N. benthamiana* and host soybean, and silencing *NbRZ-1A*/*1B*/*1C* proteins suppressed plant immunity, suggesting that RZ-1 proteins positively regulated plant resistance. In addition, complementation of GmRZ-1A or NbRZ-1A in *NbRZ-1A*-silenced leaves rescued susceptibility phenotypes, indicating that at least, RZ-1A proteins play a conserved role in immunity in diverse plants. Thus, these results extend new functions of RZ-1A in plant immunity.

In this study, PsFYVE1–NbRZ-1A interaction affected transcription and pre-mRNA AS of immune-related genes, which provides new clues for how *Phytophthora* pathogens reprogram plant immunity by changing transcriptional and post-transcriptional processes. To successfully colonize plants, pathogens produce intracellular effectors that inhibit defense responses by different molecular mechanisms, including modulating gene expression. Two nuclear effectors of *Magnaporthe oryzae*, MoHTR1 and MoHTR2, modulate disease susceptibility of rice via reprograming the transcription of many immunity-associated genes [68]. In addition to transcriptional control of gene expression, post-transcriptional processes, notably AS, emerge as key mechanisms for gene regulation during infection. For example, *P. infestans* RXLR effector SRE3 interacts directly with splicing factors U1-70K, SR30, and SR45 to manipulate AS of plant immune processes [69]. In this study, NbRZ-1A associated with the components of spliceosome complex, including GRP7, GRP8, and U1-70K, and PsFYVE1 attacked the NbRZ-1A–GRP7 interaction. The results suggest that PsFYVE1 can manipulate AS through interaction with NbRZ-1A to indirectly interfere with the spliceosome. Furthermore, RNA-seq analysis of PsFYVE1-expressing and *NbRZ-1A*-silenced leaves shed light on that PsFYVE1 and NbRZ-1A co-regulated transcription and pre-mRNA AS of many plant genes. Biochemical experiments combined with molecular-genetic analysis revealed that overexpression of PsFYVE1 and silencing of *NbRZ-1A* subverted plant defense responses by promoting transcription or AS of susceptibility factors and repressing transcription of positive regulators of plant immunity. These results provide evidence that transcriptional changes and AS regulation can co-exist simultaneously in immunomodulation, in connection with regulation properties of RNA-binding proteins.

In addition, the significant changes in different splicing variants of three genes were verified. PsFYVE1 overexpression or *NbRZ-1A* silencing induced intron retention of *Niben101Scf01240g06009*, *Niben101Scf01702g02006*, and *Niben101Scf03790g01009* (**S11 Fig**). Sequence analysis of *Niben101Scf01240g06009* and *Niben101Scf03790g01009* showed that they share similarity with genes encoding eukaryotic initiation factors in *A. thaliana*, which help stabilize the formation of the translation initiation complex to provide regulatory mechanisms in translation initiation [70, 71]. The sequence of *Niben101Scf01702g02006* is similar to that of a gene encoding a vacuole-sorting protein in *A. thaliana*, which are of great importance in autophagy and auxin transport in plants [72, 73]. These imply that PsFYVE1 and NbRZ-1A may have a fine-tuning role in developmental pathways.

In conclusion, we systematically analyzed the FYVE domain-containing protein family in oomycetes by bioinformatics, and identified functional characteristics of PsFYVE1, a FYVE protein secreted from the plant pathogen *P. sojae*. PsFYVE1 is a novel effector that interacts with plant RZ-1A proteins to manipulate the transcription and splicing process by disrupting the association of RZ-1A with splicing factor. These findings highlight a novel mechanism in which a new virulence factor of *Phytophthora* hijacks plant RZ-1A proteins to overcome plant defenses.

## Materials and methods

### Bioinformatics analyses

The genomes of organisms used in this study were retrieved from NCBI and JGI databases. To predict FYVE domain-containing proteins in each genome, the Hidden

Markov Model (HMM) profile of the FYVE domain (PF01363) was obtained from the Pfam database, followed by searching against each genome using the HMMER tool (E value cut-off <1e-5). To investigate phylogenetic relations of FYVE proteins between oomycetes and other organisms, the selected FYVE proteins were used to construct a phylogenetic tree following a neighbor-joining algorithm with 1,000 bootstrap replicates in MEGA 7 software. SignaIP 3.0 online tool was used to predict the signal peptide. Potential additional domains in each FYVE protein were predicted by searching against the Pfam database.

### Plants and microbe cultivation

*Nicotiana. benthamiana* plants were grown in a greenhouse under a 16 h day at 25°C and an 8 h night at 22°C. Four-leaf-stage *N. benthamiana* plants were used for VIGS and 6-8 weeks old *N. benthamiana* leaves were used for inoculations of *Phytophthora*. Etiolated soybean seedlings were grown without light for 5 days and then under a 16 h day at 25°C and an 8 h night at 20°C before inoculation. *Phytophthora. sojae* (P6497) and *Phytophthora. capsici* (LT263) strains were routinely maintained on 10 % vegetable (V8) juice medium at 25°C in the dark.

### Yeast signal sequence trap system

The yeast signal trap system was performed as following described [74, 75]. The signal peptide of PsFYVE1 or Avr1b (as positive control) was ligated into vector pSUC2T7M13ORI (pSUC2), which carries a truncated invertase gene (SUC2) lacking both the initiation Met and signal peptide. pSUC2 recombinant plasmids were transformed into yeast strain YTK12 using the lithium acetate method. Transformed colonies could grow on CMD-W (minus Trp) plates after 48 h of incubation at 30°C. To assay for invertase secretion, colonies were transferred to YPRAA plates. Invertase enzymatic activity was detected by the reduction of 2, 3, 5-triphenyltetrazolium chloride (TTC) to insoluble red-colored triphenylformazan. The pSUC2 empty vector was used as the negative control.

### Plasmid construction

PsFYVE1 without a signal peptide and the FYVE domain fragment was cloned from the complementary DNA (cDNA) of *P. sojae* mycelia. In the *P. sojae* transformation experiment, the FYVE domain fragment was ligated into pTOR using a one-step cloning kit (Vazyme Biotech, Nanjing, China). In the inoculation assays, the amplified PsFYVE1 fragment was ligated into pBinGFP4 (C-terminal tag green fluorescent protein (GFP) fusion) [76]. Full-length RZ-1 genes were cloned from *N. benthamiana* and soybean cDNAs. Full-length NbGRP7, NbGRP8, and NbU1-70K genes were coned from *N. benthamiana* using gene-specific primers. GmRZ-1A^syn^ and NbRZ-1A^syn^ were synthesized by Nanjing Genscript (Nanjing, China) and ligated into pCAM1300. Individual colonies of each construct were tested by PCR and verified by sequencing. The cloning primers are shown in **S5 Table.** Primers were synthesized by Sangon Biotech (Shanghai, China).

### *A. P. sojae* transformation and silencing efficiency detection

The *P. sojae* transformation was conducted as previously described [77, 78]. The silencing sequence was designed in the FYVE region of PsFYVE1. *P. sojae* strain P6497 was maintained on V8 juice agar. Before liquid culture, *P. sojae* discs were inoculated onto nutrient pea agar plates. Mycelia were harvested and then cultured in nutrient pea broth liquid medium for two days. Mycelia were washed in 0.8 M mannitol and then placed in an enzyme solution to incubate for 40-50 min at 25°C with gentle shaking. Protoplasts were harvested by centrifugation at 1,500 rpm for 3 min and particles were removed in W5 solution. After 30 min, protoplasts were collected by centrifugation at 1,500 rpm for 4 min, and adjusted concentration in MMg solution to maintain osmotic pressure of protoplasts. One milliliter of protoplasts was added 35 µg of transforming DNA and incubated for 10 min on ice. Then, three successive aliquots of 580 µL of fresh 40% polyethylene glycol (PEG) solution were pipetted into the protoplast suspension and gently mixed. After a 20-min incubation on ice, 2, 8, 10 mL of pea broth liquid medium containing 0.5 M mannitol was added to the mixed solution every 2 minutes, and protoplasts were incubated 14-16 hours to regenerate in the dark. Regenerated protoplasts were harvested by centrifugation at 2,000 rpm for 5 min, mixed thoroughly in pea broth solid medium containing 50 µg/mL G418 and 50 µg/mL ampicillin, and then poured in plates. Transformed colonies were observed after 2 days of incubation at 25°C and again covered with V8 juice medium. After 2-3 days, transformants were selected based on the transcription level of PsFYVE1. Total RNA was extracted from mycelia of transformants cultured in V8 juice medium by using an RNA-simple Total RNA Kit (Tiangen Biotech, Beijing, China). PsFYVE1 specific primers (**S5 Table**) were used to quantify expression level and evaluate silencing efficiency.

### Soybean and *N. benthamiana* inoculation assays

In soybean inoculation assays, *P. sojae* transformants were cultured in V8 juice medium without G418 selection for 5 days. Hypocotyls of etiolated soybean seedlings (Chinese susceptible cv. HF47) were inoculated with *P. sojae* transformant mycelia. Lesion lengths were measured at 48 hours post inoculation (hpi), and infection photographs were also taken at 48 hpi. Infected soybean seedlings of equal quality were collected to quantify the relative biomass of *P. sojae* by qRT-PCR. Primers of the *P. sojae* actin gene and soybean actin gene are listed in **S5 Table**. qRT-PCR reactions were performed on an ABI Prism 7500 Fast real-time PCR System (Applied Biosystems, Foster City, CA, USA).

For *P. capsici* infection on *N. benthamiana* leaves, overexpression of all proteins was confirmed by western blotting, and pBinGFP4 empty vector was used as the negative control. Leaves were detached 24 h after agroinfiltration and then inoculated with *P. capsici*. The *P. capsici*-inoculated leaves were put into a growth chamber at 25°C, and lesion areas (cm^2^) were scored at 36 hpi under UV light. Primers specific for *P. capsici* and *N. benthamiana* actin genes (**S5 Table**) were used to quantify the relative biomass of *P. capsici*.

### Protein expression and lipid filter-binding assays

*Escherichia coli* cells containing plasmids encoding each fusion protein were grown at 37°C to an OD_600_ of 0.4-0.6. Protein expression was induced with 0.1-0.5 mM IPTG at 16°C for 16-20 h. The GST fusion proteins were collected, sonicated and purified using glutathione-sepharose 4B beads (GE Healthcare, China).

Lipid filter-binding assays were conducted as previously described [13, 21]. Lipids were dissolved in a mixture of methanol: chloroform: water (20: 10: 8) and then spotted at 100 pmol onto Hybond-C extra membranes. The membranes were dried at room temperature for 1 h and then incubated in blocking buffer (50 mM Tris-HCl, 150 mM NaCl, 0.1% (v/v) Tween 20, and 2 mg/mL fatty acid-free BSA, pH 7.5) at room temperature for 1 h. Purified GST-fusion proteins, 20-30 mg, were added to the blocking buffer, and the membranes were incubated overnight at 4℃. The membranes were washed in TBST (50 mM Tris-HCl, 150 mM NaCl and 0.1% (v/v) Tween 20) 10 times over 50 min. Bound proteins were detected with mouse anti-GST antibody at a ratio of 1:2,000. After inoculation for 1 h, the membranes were washed using TBST 12 times over 1 h and incubated with a 1:5,000 dilution of the HRP-conjugated anti-mouse secondary antibody for 1 h. The membranes were washed in TBST 12 times over 1 h, and then visualized by an Enhanced Chemiluminescence (ECL) reagents according to the manufacturer’s instructions.

### Oxidative burst assay

The oxidative burst assay was carried out as previously described [79, 80], and used to determine the role of indicated construct proteins in the ROS burst in response to flg22 and chitin. A luminol-based assay was performed as following described. After protein overexpression for 48 h, leaf discs were incubated with 200 µL of water in a 96-well plate overnight in the dark at 25°C. The oxidative burst of leaf discs was detected using 200 µL of test buffer (1 µM flg22, 100 µM luminol, and 20 µg/mL horseradish peroxidase) (100 µg/mL chitin, 100 µM luminol, and 20 µg/mL L-012) and recorded by a Promega GloMax Navigator microplate luminometer (Promega Biotech, Beijing, China). GraphPad Prism software was used to analyze data.

### Western blotting

After agroinfiltration for 48 h, proteins were extracted and fractionated by SDS-PAGE. Separated proteins were transferred from the gel to a PVDF membrane and then blocked using PBST (PBS with 0.1% Tween 20) containing 5% non-fat milk at room temperature for 30-40 min. Anti-GFP (1:5,000; #M20004; Abmart Biotech, Shanghai, China) and anti-HA (1:5,000; #M20003; Abmart Biotech, Shanghai, China) antibodies were added to PBSTM (PBST with 5% non-fat dry milk) and incubated at room temperature for 180 min, followed by washes with PBST three times (5 min each). The membranes were incubated with a goat anti-mouse IRDye 800CW antibody (Odyssey, no. 926-32210; Li-Cor) at a ratio of 1:10,000 in PBSTM for 30 min at room temperature and then washed three times with PBST. Proteins were visualized by excitation at 800 nm.

### Co-immunoprecipitation assays

*A. N. benthamiana* leaves were collected 48 h after initial agroinfiltration and then frozen in liquid nitrogen and ground to powder using mortar and pestle. The total proteins were extracted using RIPA lysis buffer (50 mM Tris pH 7.4, 150 mM NaCl, 1% Triton X-100, 1% sodium deoxycholate and 0.1% SDS) and centrifuged at 4°C for 10 min at 12,000 × g. The supernatant was transferred to a new tube, and appropriate SDS-PAGE sample loading buffer was added into tubes to detect proteins by western blotting. For GFP-IP, 1-4 mL of supernatant was incubated at 4°C with GFP-Trap_A beads (Chromotek, Planegg-Martinsried, Germany) overnight. Beads were collected by centrifugation at 2,500 × g and then washed three times in 1-4 mL of washing buffer (10 mM Tris-Cl, 150 mM NaCl, and 0.5 mM EDTA, pH 7.5). Appropriate SDS-PAGE sample loading buffer was added to bound proteins and boiled for 8-10 min. The results were detected by western blotting using anti-GFP and anti-HA antibodies. The images were caught by Odyssey CLx System.

### VIGS of NbRZ-1 genes in *N. benthamiana*

Tobacco Rattle Virus (TRV)-based VIGS system was used to silence *NbRZ-1A*, *NbRZ-1B*, and *NbRZ-1C* gene in *N. benthamiana*. Fragments from each gene were cloned into the pTRV2 vector. The TRV:GUS vector was used as the control. Four-leaf stage *N. benthamiana* plants were used in agroinfiltration containing a mixture of pTRV1 and pTRV2 vectors at OD_600_ of 0.5 each. Silenced leaves were inoculated, and silencing efficiency was detected at 30 days post-infiltration (dpi) of pTRV vectors.

### Confocal microscopy and bimolecular fluorescence complementation

The *N. benthamiana* leaves were cut and mounted in water and analyzed using an LSM 710 laser scanning microscope with a × 63 objective lens (Carl Zeiss, Jena, Germany). *RZ-1* genes and *PsFYVE1* (without signal peptide) sequences were cloned into YFPn and YFPc vectors separately. The *A. tumefaciens* strains GV3101 harboring indicated plasmids were mixed at final OD_600_ of 0.5 each, and 6 weeks old *N. benthamiana* plants were used for infiltration. Co-locational fluorescence was observed 36-48 hpi using confocal microscopy.

### Luciferase complementation assays

Lluciferase complementation assays were carried out as previously described [69, 81]. *PsFYVE1*, *PsFYVE1* mutant genes (without a signal peptide), and *N. benthamiana GRP7*, *GRP8*, and *U1-70K* genes were inserted into pCAMBIA1300-nLUC (nLUC), whereas soybean *RZ-1A*/*1B*/*1C* genes and *N. benthamiana RZ-1A*/*1B*/*1C* genes were inserted into pCAMBIA1300-cLUC (cLUC).

*A. tumefaciens* strains GV3101 harboring indicated plasmids were mixed at final OD_600_ of 0.5 each and then infiltrated into *N. benthamiana* leaves. The *N. benthamiana* leaves were collected at 48 h after infiltration and then incubated with 1 mM luciferin substrate (catalog no. N1110; Promega Biotech, Beijing, China). LUC images were captured with a Tanon-5200 Multi Chemiluminescent Imaging System (Tanon, China). The *N. benthamiana* leaf discs were incubated with 1 mM luciferin substrate in a 96-well plate, and activity of luciferase reporter gene (LUC) was detected using Promega GloMax ® Navigator microplate luminometer.

## Acknowledgments

We appreciate Dr. Suomeng Dong at Nanjing agricultural University and Dr. Jie Huang at Nanjing agricultural University for their valuable suggestions.

## Author Contributions

D.D. and D.S. conceived and designed the experiments. X.L., Z.Y., W.S., J.S., Z.Y., and M.J. performed the experiments and/or data analyses. X.L., D.S., and D.D. wrote and modified the manuscript. All authors read and approved the final manuscript.

## Supporting information

**S1 Fig. Phylogenetic analysis of FYVE domain-containing proteins.** The FYVE domain-containing protein sequences derived from *P. sojae*, *A. thaliana*, *S. cerevisiae*, and *H. sapiens* were used to constructed the phylogenetic tree

**S2 Fig. PsFYVE1 contains classic FYVE domain and shows high expression during infection.** (A) Multiple sequence alignment of PsFYVE1 and its orthologs. Residues involved in PI3P are indicated above the alignment. Other residues at conserved hydrophobic positions are shown in green type. (B) Expression of PsFYVE1 during *P. sojae* growth and infection of soybean. Total RNA was extracted from mycelia (MY) or soybean root tissues infected with zoospores of *P. sojae* P6497 at 3, 6, 12, and 24 hpi. Transcript levels of PsFYVE1 were determined by qRT–PCR.

**S3 Fig. PsFYVE1 carries a functional secretory signal peptide in yeast invertase secretion assay.** The 24 amino acid of *PsFYVE1* N-terminal signal peptide sequence was cloned into the yeast vector pSUC2. The predicted signal peptide sequence of PsAvr1b was used as a positive control and the empty vector was used as a negative control. CMD-W (minus Trp) plates were used to select yeast strain YTK12 carrying the pSUC2 vector. YPRAA media was used to indicate invertase secretion. An enzymatic activity test based on the conversion of TTC to the insoluble red-colored product was used to confirm invertase secretion.

**S4 Fig. Silencing of *PsFYVE1* does not affect mycelial growth.** (A) Growth rate of *PsFYVE1*-silenced transformants. Wild-type (WT), *PsFYVE1*-silenced transformants (T5/ 8/ 15) and T9 (a non-silenced transgenic line used as a negative control) were growing on V8 medium for 5 days. Experiments were repeated three times with similar results. Scale bar represents 1 cm. (B) Silencing efficiency of transformants. Relative transcript levels of *PsFYVE1* gene in the different transformants were determined by qRT-PCR using soybean actin gene as a reference gene. (C) Growth rate of the different *PsFYVE1-*silenced transformants. The colony growth rate was calculated based on the diameters of colonies. All data are shown as mean ± _SD_ (n = 3-8). Asterisks indicate statistical significance: **P<0.01, two-sided unpaired *t* test.

**S5 Fig. Phsphoinositide-binding activities of PsFYVE1 and its FYVE domain.** (A) PI3P binding site residues in FYVE domains of EEA1, Vps27p, and PsFYVE1. Multiple alignment of FYVE finger sequences of three FYVE proteins and PsFYVE1^AAAA^ mutant are shown. Residues involved in PI3P are indicated above the alignment. Other residues at conserved hydrophobic positions are shown in green type. The amino acid mutations of PsFYVE1^AAAA^ are indicated below the alignment by arrows. (B) Phsphoinositide-binding activities of PsFYVE1 and FYVE domain of PsFYVE1 proteins. The Hybond-C membranes carried individual phosphoinositides, including PI, PI3P, PI4P, PI5P, PI4,5P_2_ and PI3,4,5P_3_ (100 pmol per spot) were incubated with the indicated recombinant proteins overnight at 4 ℃. GST-labelled a mouse FYVE protein, GST-EEA1, was used as a positive control and the GST empty vector was used as a negative control. PsFYVE1^AAAA^ was a PI3P-binding sites mutant of PsFYVE1 and GST-2×FYVE was tandem repeat FYVE domain of PsFYVE1 fused GST tag. The bound proteins were detected with anti-GST monoclonal antibodies followed by an HRP-conjugated secondary antibody.

**S6 Fig. PsFYVE1 interacts with NbRZ-1A, NbRZ-1B, and NbRZ-1C**. (A) Co-IP of PsFYVE1 and NbRZ-1A/1B/1C. IP of protein extracts from agroinfiltrated leaves using GFP-Trap confirmed that PsFYVE1-HA associated with GFP-tagged NbRZ-1A/1B/1C in *N. benthamiana*, but not the GFP control. The positions of the expected protein bands are indicated by asterisks. Protein sizes are indicated in kDa, and protein loading is visualized by Ponceau stain. (B) Interaction and co-localization of PsFYVE1 and NbRZ-1A/1B/1C in BiFC assay. *A. tumefaciens* cells harboring YFPn (Yn) and those harboring YFPc (Yc) were co-infiltrated into *N. benthamiana*. The fluorescent signals were detected by confocal laser microscopy at 48 hours after infiltration and DAPI was used to a nucleus staining. Scale bar represented 20 μm.

**S7 Fig. Transcription profiles and sequence analysis of RZ-1A proteins**. (A) Transcription levels of *RZ-1A*/*1B*/*1C* in RNA-seq data. The data were obtained from the infected soybean roots (cultivar: Williams 82) by *P. sojae* (P6497) zoospores and the infected *N. benthamiana* leaves by *P. parasitica* zoospores. The gene expression levels were normalized as reads per kilo-base per million (RPKM) in previous reports [54, 55]. (B) Sequence alignment of GmRZ-1A and NbRZ-1A. BioEdit was used for the multiple alignments of the sequence. The positions of G-rich regions are marked with lines below the sequence.

**S8 Fig. Complementation of GmRZ-1A^syn^ and NbRZ-1A^syn^ recovers the TRV:NbRZ-1A phenotype.** (A) Complementation of GmRZ-1A^syn^ or NbRZ-1A^syn^ on the NbRZ-1A-silenced *N. benthamiana* leaves. *N. benthamiana* leaves were infiltrated with synthetic GmRZ-1A^syn^ or NbRZ-1A^syn^ construct at 30 days after infiltration of TRV vectors. The photographs were taken at 36 hpi under UV light. Dashed lines show the lesion areas. (B) Immunoblot analyses of indicated recombinant proteins. Total proteins were extracted at 48 hours after infiltration. Protein expression was confirmed by western blotting using anti-GFP antibody. The positions of the expected protein bands are indicated by asterisks. Protein size in kDa is shown, and protein loading is visualized by Ponceau stain. (C) Average lesion size of inoculated leaves. Lesion size (cm^2^) was measured at 36 hpi. Error bars represent the mean ± _SD_ (n = 12). Asterisks indicate statistical significance: **P<0.01, two-sided unpaired *t* test. (D) Complementation of GmRZ-1A^syn^ or NbRZ-1A^syn^ on flg22-induced ROS burst tests. Flg22-induced ROS were detected on the leaves at 48 hours after infiltration. Relative luminescence units (RLU) indicate the relative amounts of H_2_O_2_ production induced by 1 μM flg22. Means and standard deviations from six replicates are shown.

**S9 Fig. RZ-1A C-terminus is essential for interaction with PsFYVE1 and immune-function.** (A) Domain architecture of GmRZ-1A, NbRZ-1A and their mutants. Schematic drawings of a series of truncated mutants including GmRZ-1A^ΔCT^ (1-110 amino acid), GmRZ-1A^ΔRRM^ (80-208 aa), NbRZ-1A^ΔCT^ (1-110 aa), and NbRZ-1A^ΔRRM^ (78-209 aa) are shown. The percentage of glycine (G), arginine (R), and aspartic acid (D) residues of the C-terminus included the zinc finger are shown in parentheses. RRM, RNA recognition motif, Zf, zinc finger, G-rich, glycine-rich. (B) The interaction between GmRZ-1A/NbRZ-1A mutants and PsFYVE1 in split-LUC assays. The leaves were photographed using Tanon-5200 Multi under chemiluminescent imaging system, and 16 leaf discs were used to measure the luminescence 48 h after co-expression of the indicated proteins. (C) *P. capsici* infection assay on *N. benthamiana* leaves. GmRZ-1A, NbRZ-1A and their corresponding mutants (GmRZ-1A^ΔCT^ and NbRZ-1A^ΔCT^) were expressed in leaves. *P. capsici* mycelia were inoculated on the infiltrated leaves at 24 h after Agro-infiltration. The photographs were taken at 36 hpi under UV light. Dashed lines showed the lesion areas. (D) Average lesion size of inoculated leaves. Lesion size (cm^2^) was measured at 36 hpi. Data are shown as mean ± _SD_ (n = 12). Asterisks indicate statistical significance: **P<0.01, two-sided unpaired *t* test. (E) Immunoblot analyses of the indicated recombinant proteins. Total proteins were extracted at 48 hours after infiltration. Protein expression was confirmed by western blotting using an anti-GFP antibody. The positions of the expected protein bands are indicated by asterisks. Protein sizes are indicated in kDa, and protein loading is visualized by Ponceau stain. (F) Nuclear speckle localization of RZ-1As is in a C-terminus-dependent manner. The confocal images of *N. benthamiana* epidermal cells transiently expressing GmRZ-1A-GFP, NbRZ-1A-GFP, GmRZ-1A^ΔCT^-GFP, and NbRZ-1A^ΔCT^-GFP were photographed at 48 hours after infiltration. Scale bar represented 20 μm.

**S10 Fig. Both PsFYVE1 and NbRZ-1A regulate transcription of plant immunity-related genes**. (A) Venn diagram indicating the overlap of differentially expressed genes (DEGs) in PsFYVE1-expressing and *NbRZ-1A*-silenced *N. benthamiana* leaves at 36 hpi. Among two groups of DEGs, 60 and 19 genes were significantly upregulated (red face) or downregulated (blue face) simultaneously. (B) Gene ontology (GO) enrichment for biological process domains of the 79 DEGs simultaneously present in both treatments. (C-E) Relative expressional levels of *NbHCT* (B), *NbEIN2* (C), and *NbSUS4* (D). Three defense related DEGs including NbHCT, NbEIN2, and NbSUS4 for validation of transcriptional changes by qRT-PCR analysis. Total RNA of the indicated *N. benthamiana* leaves was extracted for qRT-PCR assay using overexpression of GFP and TRV:GUS as controls. Means and standard deviations from three replicates are shown. (F) *P. capsici* infection assay on the *NbHCT*-, *NbEIN2*-, and *NbSUS4*-silenced *N. benthamiana* leaves. *P. capsici* mycelia were inoculated on the leaves 30 days after Agro-infiltration. The photographs were taken at 36 hpi under UV light. Dashed lines show the lesion areas. Lesion sizes (cm^2^) of infected *N. benthamiana* leaves are measured at 36 hpi. Data are shown as mean ± _SD_ (n = 9). Asterisks indicate statistical significance: **P<0.01; * P<0.05, two-sided unpaired *t* test.

**S11 Fig. Both PsFYVE1 and NbRZ-1A regulate pre-mRNA alternative splicing**. (A-C) Validation of splicing ratios of *Niben101Scf01240g06009* (A), *Niben101Scf01702g02006* (B), and *Niben101Scf03790g01009* (C) genes by qRT-PCR. The schematic shows models of two isoforms of three genes (upper panel), in which the asterisks indicate a premature termination codon. qRT-PCR was performed using isoform-specific primers to measure the splicing ratios of three genes (lower panel). Total RNA was extracted from the indicated *N. benthamiana* leaf tissues infected with *P. capsici* at 36 hpi, including overexpression of GFP (a control), PsFYVE1-GFP, TRV:GUS (a control), and *NbRZ-1A*-silenced (TRV:NbRZ-1A). Means and standard deviations from three replicates are shown.

**S1 Table. Putative targets of PsFYVE1 by immunoprecipitation assay.**

**S2 Table. Putative targets of NbRZ-1A by immunoprecipitation assay.**

**S3 Table. Differentially expressed genes identified in both PsFYVE1-overexpressing and NbRZ-1A-silenced samples at 36 hpi.**

**S4 Table. A list of changed alternative splicing events. S5 Table. Primers used in this study.**

